# Buffer specificity of ionizable lipid nanoparticle transfection efficiency and bulk phase transition

**DOI:** 10.1101/2025.01.17.633509

**Authors:** Cristina Carucci, Julian Philipp, Judith A. Müller, Akhil Sudarsan, Ekaterina Kostyurina, Clement E. Blanchet, Nadine Schwierz, Drew F. Parsons, Andrea Salis, Joachim O. Rädler

**Author notes:** Corresponding Authors Prof. Joachim Radler, Prof. Drew Parsons. CC and JP contributed equally to the work.

## Abstract

Lipid nanoparticles (LNPs) are efficient and safe carriers for mRNA vaccines based on advanced ionizable lipids. It is understood that the pH dependent structural transition of the mesoscopic LNP core phase plays a key role in mRNA transfer. However, buffer specific variations in transfection efficiency remain obscure. Here we analyze the effect of buffer type on the structure and transfection efficiency of LNPs. We find that LNPs formulated with the cationic ionizable lipid DLin-MC3-DMA (MC3) in citrate compared to phosphate and acetate buffers exhibit earlier onset and stronger mRNA-GFP expression in-vitro. Using synchrotron small angle X-ray scattering (SAXS) we determine the pH dependent mesophases of ionizable lipid MC3/cholesterol/water dialyzed against the various buffers. The results show that the phase transition with decreasing pH from inverse micellar cubic to inverse hexagonal (Fd3m-H_II_) is shifted by one unit to a lower transition pH for acetate and phosphate compared to citrate buffer. Based on continuum theory and ion specific adsorption obtained from all-atom MD simulations, we propose a mechanism for buffer specificity. Citrate stabilizes the inverse hexagonal phase thus shifting the formation of H_II_ to a higher pH. By contrast, phosphate and acetate stabilize L_II_. We propose that the Fd3m-to-H_II_ transition, which is facilitated in citrate buffer, is responsible for a sensitized pH-response of the LNP core phase. This, in turn, enhances endosomal release efficiency and accounts for the earlier onset of gene expression observed in LNPs prepared with citrate buffer.

## Main

Gene delivery carriers have been developed and improved for decades but reached an unprecedented breakthrough through successful, safe and efficient delivery of mRNA-based vaccination during the COVID-19 pandemic.^1–3^ Specifically, lipid nanoparticle (LNP) formulations have been approved by the Food and Drug Administration (FDA) due to favorable properties such as colloidal stability, low toxicity, and controlled size. While early cationic lipoplexes proved toxic to human cells, LNPs based on ionizable lipids exhibit pH dependent lipid headgroup ionization and hence less toxic surface charge. Furthermore, once endocytosed, LNPs overcome endosomal entrapment via endosomal fusion in a pH dependent manner. LNPs are internalized at physiological pH and experience protonation during the time course of the early endosomal maturation with pH values decreasing to (6.5-5.0).^4,5^ The process of charging ionizable lipid headgroups facilitates endosomal fusion, releasing mRNA into the cytosol. The exact process of how LNPs fuse with the endosome membrane is the subject of intense research. In cationic lipid based lipofection it has been rationalized that the mesoscopic bulk structure of lipoplexes affects fusogenicity, with the inverse hexagonal phase^6^ and cubic phase^7^ being particularly fusogenic compared to lamellar internal packing. With the advent of ionizable lipids, the LNP core mesostructures become pH-dependent and multiple phases and structural transitions emerge as a function of decreasing pH.^8–10^ It is generally understood that lyotropic mesophases depend on the lipid chain splay described by the Israelachvili shape factor.^11,12^ Ionizable lipids with strongly conic shape form inverted phases, which with increasing headgroups size range from inverse micellar disordered, L_II_, to disconnected inverse micellar cubic, I_II_, to inverse hexagonal, H_II_ and bicontinuous cubic phases, Q_II_. Thereby the (I_II_-H_II_) as well as (H_II_-Q_II_) transitions change connectivity of the lipid network.^13^ In recent work, we showed that specifically the ionizable lipid MC3 shows a structural phase transition with decreasing pH from inverse micellar cubic phase with space group Fd3m to inverse hexagonal (denoted Fd3m-H_II_ transition in the following).^14^ X-ray scattering from full LNPs also provide evidence that a similar structural transition occurs within the LNPs as a function of pH, assuming that the LNP core mesophase is predominantly formed by ionizable excess lipid and cholesterol. The pH-dependent lipid core transition has been suggested as a critical factor in inducing endosomal fusion and hence mRNA escape efficiency.^14^ Therefore, further investigation of ionizable lipid/cholesterol bulk phases as LNP core mimics is valuable for gaining insight into the LNP-endosomal fusion mechanism.

Here, the critical pH-value of the structural transition coincides with the pK*_a_* value of the ionizable lipid. The regulation of pH in both chemical and biological systems is carried out by buffers, a mixture of a weak acid (HA) or base (B) with its conjugate base (A^-^) or acid (BH^+^). According to the Henderson-Hasselbalch equation, the only important parameter for the choice of buffer is the pK_a_ and the concentrations of the buffer components. Little attention has been paid to the chemical identity of the weak electrolytes and their conjugate species used to prepare the buffer.^15^ However, starting from the pioneering work by Ninham and coworkers on restriction enzyme activities,^16^ several studies have shown that the chemical nature of the buffer, even at the same nominal pH, can have important unexpected effects on the investigated biosystem. For example, buffers have been found to affect specifically the behavior of proteins, including lysozyme electrophoretic mobility^17^ and adsorption,^18^ Brownian motion of BSA,^19^ and more recently, DNA thermal stability,^20^ DNA interactions with lipid bilayers,^21^ and the formation of a protein corona around nanoparticles.^22^ In fact, “specific buffer effects” can be included in the wider classification of “ion specific effects” first observed by Hofmeister in 1888.^23^ The “Hofmeister series” is an order based on ion induced protein precipitation (salting out) or solubilization (salting in). A conventional explanation of the Hofmeister series was proposed ^24,25^ invoking Jones and Dole’s work ^26^ on the viscosity of aqueous salt solutions. Ions were classified as “kosmotropic” (order maker) or chaotropic (disorder maker) on the basis of their interaction with water quantified through the value (and the sign) of the Jones-Dole viscosity B coefficient. More recent theoretical and simulation work explains ion specificity as the result of a delicate interplay between electrostatic, hydration and ion-dispersion forces.^27,28^ Whatever the details of the mechanisms explaining “Hofmeister phenomena”, it must be considered that ions play an important role to modulate biological mechanisms in a way which is not fully understood. For example, Meulewaeter et al.^29^ found that Tris and HEPES buffers improved cryoprotection and, more importantly, the transfection efficiency of mRNA-LNPs compared to PBS buffer. What emerges from previous studies is that the mechanism of transfection, based on bulk phase transitions, is controlled not only by pH but is also specifically affected by the buffer ions used to control pH.^30,31^

The H^+^ charge equilibrium of the ionizable lipid is influenced by the buffer in two ways. Interactions between buffer ions and H^+^ ions ultimately result in a non-negligible activity coefficient γ_H_ for H^+^ (and also for the lipid charge site). However, activity coefficients are only expected to be significant at very high ion and buffer concentrations, stronger than the buffer concentrations considered here. The second mechanism of the buffer effect enters through the surface potential at the lipid charge site. A term for the surface potential ψ_0_ appears in the mass balance equation relating the equilibrium constant to ion concentrations,^32^

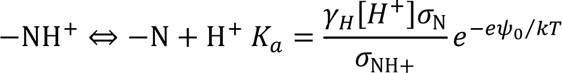

where σ_NH+_ and σ_N_ are surface densities of the protonated and deprotonated MC3 charge sites, respectively. Algebraically the term in ψ_0_ could be shifted over to *K_a_* to form an “effective “p*K_a_*” that depends on conditions, and this is sometimes done in the literature to simplify the mass action equation as an equation of ion concentrations rather than ion chemical potentials. It is misleading however and obfuscates the role of the surface potential in determining the charge of the ionizable lipid. The surface potential may be influenced quite strongly by the buffer solution, for instance via specific adsorption of buffer anions, a point which we come back to later when discussing mechanisms determining the observed specific buffer effects.

The present work aims to investigate the effect of the choice of preparation buffer solution (here, citrate, phosphate and acetate buffers) on the transfection efficiency of LNPs formed with the ionizable lipid DLin-MC3-DMA (MC3), and on the transition of bulk phases composed of only MC3/cholesterol. This reduction to only two lipids is possible since both DSPC and PEG are mostly absent in the LNP core.^9^ In order to explain the observed buffer specific transfection efficiency, we investigated bulk phase transitions at different pH and ionic strength with different buffers, through synchrotron small angle X-ray scattering (SAXS). The nearest neighbor distance, d_NN_, values, (the distance from one micelle water-core to the next), was also evaluated as a function of pH. We propose a mechanism for the observed buffer specific pH shift of the (Fd3m-H_II_) bulk phase transition in terms of changes in the area per lipid headgroup affected by pH and by ion specific adsorption based on all-atom MD simulations. This study highlights the role of buffer ions on the pH-dependent structural transitions in ionizable lipid bulk phases. We hypothesize that a pH-dependent structural transition also occurs in the excess lipid region within the LNP core phase that is critical for endosomal release. As citrate buffer facilitates the transition this leads to an earlier gene expression onset compared to phosphate and acetate buffer. The proposed structure-activity relation highlights the importance that the suitable choice of the buffer has for mRNA transfection and is therefore significant for the full development of LNP technology in nanomedicine.

## RESULTS AND DISCUSSION

### Specific buffer effects on mRNA LNPs and transfection efficiency

LNPs are composed of an external layer composed of PEG polymer with DSPC and MC3 lipids, with an internal bulk phase composed of Dlin-MC3-DMA and cholesterol. MC3 is an ionizable lipid with a p*K_a_* of 6.44,^33^ which makes it pH sensitive (Fig. 1A-B). The LNPs are internalized inside living cells at pH ∼ 7 until reaching the endosome, where mRNA is released at pH ∼ 6-6.5 (Fig. 1E). To study the buffer effects three common buffers were chosen (Fig. 1C-D).

**Figure 1.**
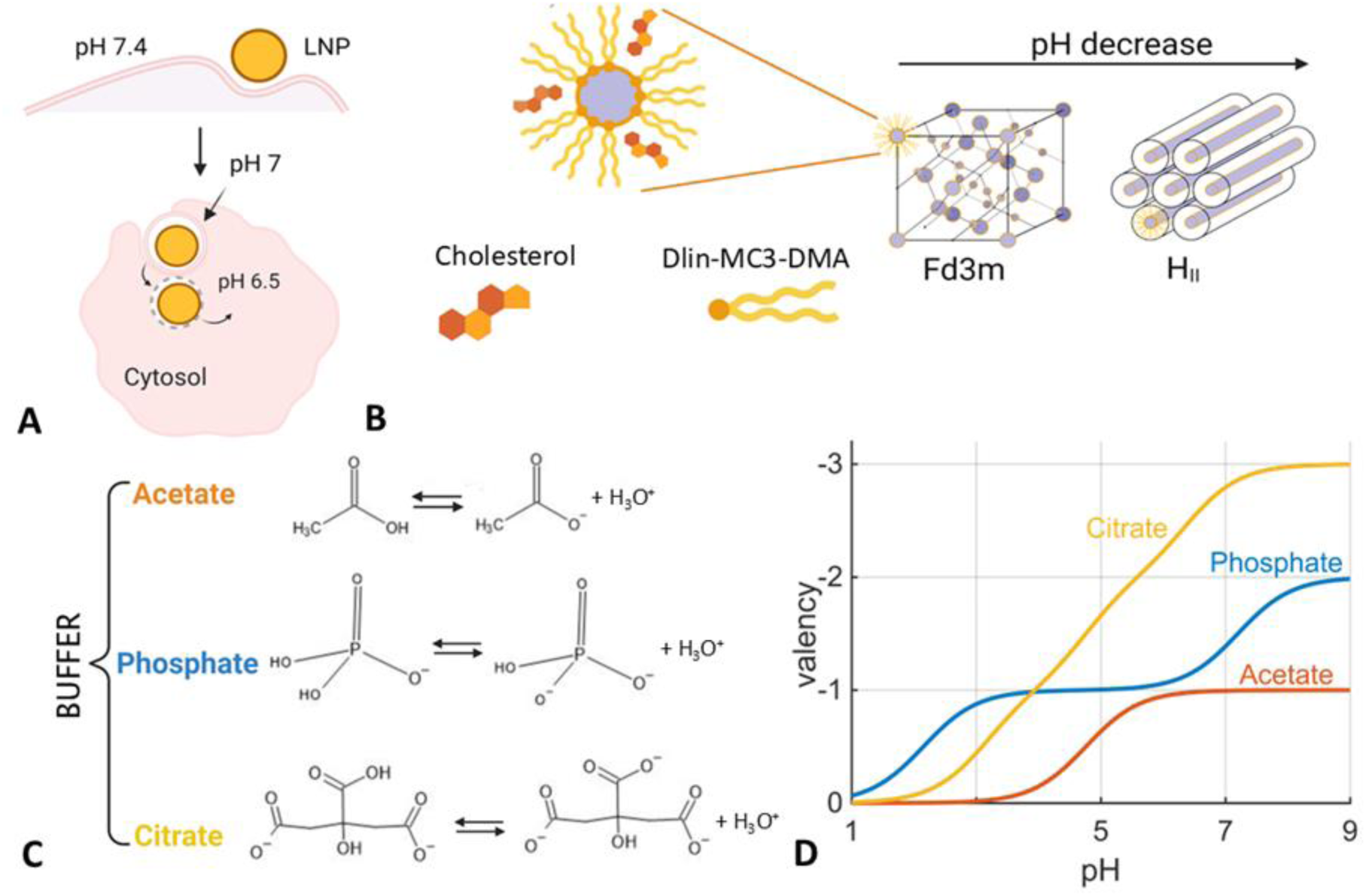
Buffer specificity of LNP mediated mRNA delivery. (A) Schematic drawing of internalization via endocytosis. Endosomal release of LNPs occurs as a consequence of acidification inside the endosome from pH 7 to about pH 6.5. Schematic drawing of bulk phase made by ionizable lipid Dlin-MC3-DMA and cholesterol together with bulk phase transition from Fd3m to H_II_ as pH decreases. (C) The buffer used during the dialysis process is chosen among a range of buffers, citrate (pK*_a_* 6.4), phosphate (pK*_a_* 7.2), and acetate (pK*_a_* 4.8). To evaluate the phase transition a wide range of pH values from 3.5 to 7.0 in a 0.5 pH unit has been studied. (D) Buffer valency at each pH depending on pK*_a_* values.

According to the Henderson-Hasselbalch equation (pH = p*K_a_* ± 1), citrate, phosphate and acetate with p*K_a_* of 6.40, 7.22 and 4.80, respectively, cover the whole pH range of the LNP route from internalization (pH 7) to mRNA release (pH 6). Sodium citrate buffer has been used in cationic lipid design for siRNA delivery,^1^ and is by far the buffer used most for LNP preparation at acidic pH. Sodium phosphate buffer was used with KCl and NaCl to store Pfizer’s mRNA vaccine.^30^ Sodium acetate buffer has been used to dilute mRNA for LNP encapsulation^34^ and as dialysis buffer for ethanol removal after LNP preparation.^35^ To investigate buffer specific effects on transfection kinetics, we prepared eGFP-mRNA LNPs in citrate, phosphate or acetate buffer as described previously,^14^ followed by dialysis into water. We characterized LNP quality by measuring size and eGFP-mRNA encapsulation. The data, presented in the supplementary information, show no buffer-specific dependency (Fig. S1 and Table S1). To access time-resolved protein expression, we performed live-cell imaging on single cell arrays (LISCA).^36^ After pre-incubation in cell culture medium supplemented with serum, LNPs in medium were transfected into the human liver carcinoma cell line (HuH7) seeded on a single-cell slide (Fig. 2A).

**Figure 2.**
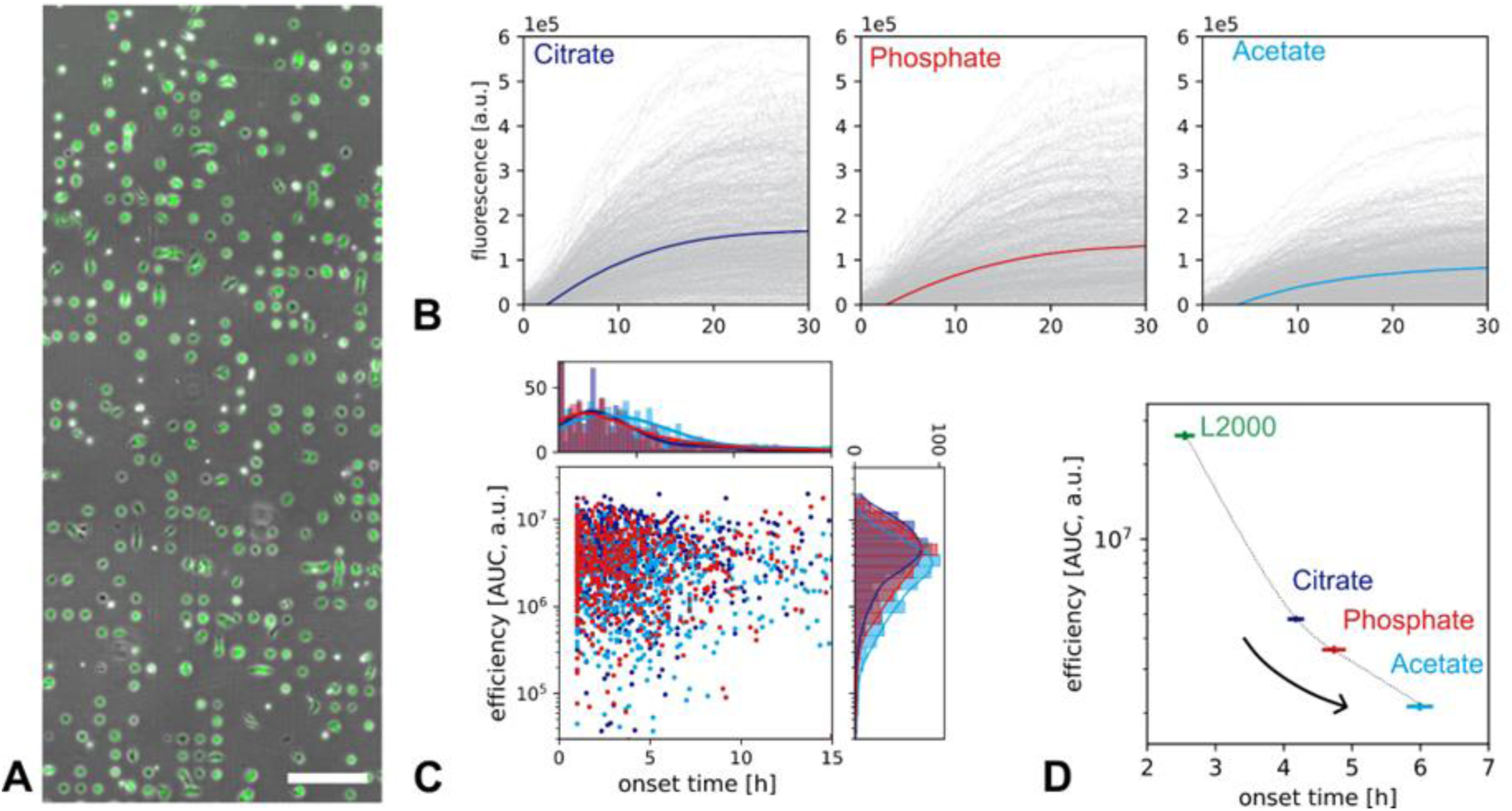
Single cell transfection experiments: (A) Culturing cells on microfabricated single-cell arrays allows recording of hundreds of single-cell fluorescence trajectories in parallel. (B) Single cell expression time courses of eGFP-mRNA/LNPs (gray lines) show different fluorescence kinetics for different preparation buffers (averaged time courses shown in color). (C) Scatter plot of single cell expression efficiencies (area under the curve) versus expression onset times (D) Mean values of single cell efficiencies (C) for the three LNP buffer conditions exhibit an apparent Hofmeister ordering. Data for lipofectamine transfection are shown as reference. In general, faster onset is correlated with higher protein expression. Error bars indicate standard error of the mean.

Observation of GFP expression kinetics of single cells resulted in a fluorescence trajectory for each cell (Fig. 2B). Distinct differences between the buffers were apparent from these traces. Averaging all traces revealed the highest overall GFP level for LNPs prepared in citrate buffer followed by those prepared in phosphate buffer and the lowest expression from the acetate LNPs. Single-cell resolution allowed for the calculation of the total protein amount per cell, expressed as the area under the curve (AUC) and the distinct onset of protein expression for every single cell (Fig. 2C). The highest mean AUC confirmed the initial observations from the fluorescence trajectories: protein expression varied depending on the buffer in which the LNP was prepared, with the highest eGFP expression for the citrate, followed by phosphate and lowest for the acetate buffer. Hence the buffer specific effect appears to follow a conventional Hofmeister series, with citrate > phosphate > acetate. The observed ordering in the onset times exhibits an inverse relation of expression efficiency and expression onset time (Fig. 2D). This confirms a correlation between fast onset and high protein expression levels as previously reported in time-resolved studies.^37^ The reference transfection with Lipofectamine^TM^ 2000 (Thermo Fisher Scientific) is consistent with this finding (Fig. 2D). These findings indicate that the underlying endosomal release mechanism is dependent on the LNP preparation buffer.

### Specific buffer effects on bulk phase transitions

Next, we investigate the impact of buffers on the phase behavior of ionizable lipid/cholesterol bulk phases as a function of pH. These two-component bulk phases serve as a model for the inner core structure of LNPs, given that PEG and DSPC are restricted to the outer shell of the LNPs. As described in the methods section, the preparation of bulk phase samples consists of three dialysis steps to mimic the LNP production by replacing the ethanol with buffer. The buffer (citrate, phosphate or acetate) at the pH of choice comes into play in the third dialysis step. Figure 3A presents the SAXS scattering profiles for MC3/cholesterol/buffer phases in the presence of citrate, phosphate, and acetate buffers. In the following sections, we describe the pH-dependent structural transitions observed. As shown in Figure 3B decreasing the pH induces a phase transition from inverse micelles (L_II_)^38^ to inverse hexagonal phase (H_II_).^38^ The inverse micellar phase undergoes ordering transitions from a disorder L_II_ through close-packed P6_3_/mcc^38,39^ towards an inverse cubic Fd3m phase.^40^ This order of inverse phases as a function of pH has been reported before for MC3/cholesterol in citrate buffer.^10^ Here we find that the same behavior is found also in phosphate and acetate buffer but with slightly shifted pH values of transitions. In citrate buffer P6_3_/mmc and Fd3m phases are formed at pH 7.0 and pH 6.5 respectively with decreasing pH.

**Figure 3.**
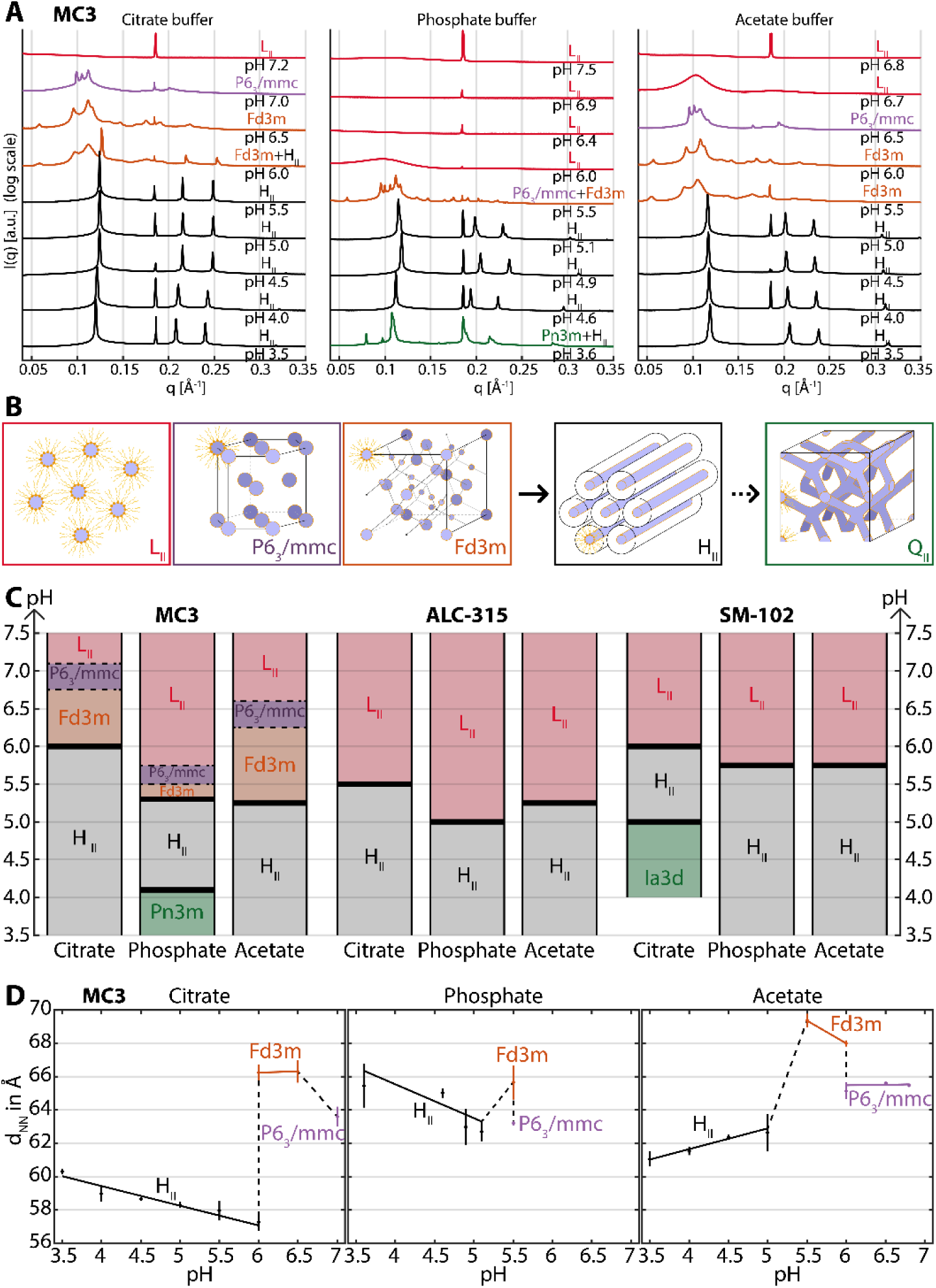
Buffer specific effects on ionizable lipid mesophases (A) SAXS measurements of MC3-cholesterol bulk samples dialyzed in the presence of 50 mM citrate, acetate and phosphate buffer and NaCl 150 mM across a pH range of approximately 3.5 to 7.5. (B) Sequence of lipid phase symmetries with increasing protonation showing the order L_II_, P6_3_/mcc, Fd3m, H_II_ and Pn3m.^38^ (C) Phase diagrams showing the buffer-specific pH dependence for MC3, ALC-315 and SM-102. (D) Nearest neighbor distance d_NN_ between centers of encapsulated water for MC3 with H_II,_ Fd3m and P6_3_/mcc phases as a function of pH for the three different buffers.

The same transitions are observed in the presence of acetate, but at a lower pH (P6_3_/mmc at pH 6.5 and Fd3m at pH 6.0). A bigger difference is observed for phosphate resulting in the L_II_ phase being found in the region from pH 7.5 to 6.0 and a mixed phase P6_3_/mmc+Fd3m at pH 5.5. The main topological transition, that is thought to be structurally disruptive and hence might correlate with endosomal fusion, occurs at the inverse cubic Fd3m to inverse hexagonal H_II_ transition point.^10^ This inverse cubic-to-inverse hexagonal (Fd3m - H_II_) transition occurs at pH*= 6.0 for citrate and at pH*= 5.0 for acetate and phosphate (Fig. 3 A-C). Remarkably the critical pH* value is shifted by one pH unit for acetate and phosphate compared to citrate. To demonstrate that the transition from inverse micellar (L_II_) to inverse hexagonal phase (H_II_) is a universal characteristic of clinically relevant ionizable lipid we also present the phase behavior of ALC-315/cholesterol and SM-103/cholesterol for all three buffers. We observe that the specific buffer effect on the L_II_-H_II_ transition follows the trend pH*(citrate)>pH*(phosphate) = pH*(acetate). This behavior can be explained by the common conic lipid shape factor which promotes inverse micellar phases at neutral pH. However, the charging of the lipid head groups, and consequently the geometric packing parameter of the lipids, is influenced by pH, temperature and salt concentration. In the following we examine these parameters in detail for the MC3 lipid.

### Nearest neighbor distance (d_NN_)

The dimensions of the bulk phase are characterized by the nearest neighbor distance (d_NN_), determined from SAXS measurements. d_NN_ is defined as the smallest distance between neighboring water cores or channels. Significant differences in d_NN_ as a function of pH are seen in Fig. 3D. d_NN_ decreased up to 13% from Fd3m to H_II_ phase in the case of citrate. For acetate a smaller difference of 9% is shown when the phase changes from Fd3m to H_II_. For phosphate the difference is even less evident, reaching up to 6% of decrease from Fd3m to H_II_ phase. d_NN_ of P6_3_/mmc decreased by about 5% when compared to the Fd3m phase for all buffers (Fig. 3D).

With the thickness of the nonpolar layer determined largely by the length of the MC3 lipid tails, a decrease in d_NN_ can be understood as a decrease in the diameter of the aqueous cores. Among all buffers, the Fd3m phase of acetate showed the highest d_NN_ of 70 Å indicating the highest water content in the bulk phase when bulk phase is dialyzed against acetate. d_NN_ decreased linearly with the increasing pH for H_II_ phase in the case of citrate and phosphate buffer due to electrostatic interactions among charged headgroups which favors a smaller d_NN_. Curiously, the d_NN_ of acetate in H_II_ phase increased with pH, showing a reverse slope compared to the other two buffers.

### Effect of ionic strength and temperature on bulk phase

The previous experiments were carried out by using the same concentration of 50 mM for all buffers at the different investigated pH values. But the charge (z) and the concentration (c) ratio between the acidic and the basic component of the buffer depends on pH (Fig. 1 C-D). For instance, at pH 5.5, say, the ionic strength for the different 50 mM buffers is 151 mM for citrate, 52 mM for phosphate, and 42 mM for acetate. To understand if the observed buffer specificity (Fig. 2) could be ascribed to an ionic strength effect of the different 50 mM buffers, SAXS measurements were performed of the LNP bulk phases dialyzed in the presence of citrate and acetate at pH 5.5 and at the same ionic strengths (namely 150, 200, and 300 mM). pH 5.5 was chosen since all buffers showed a clearly ordered structure (Fd3m or H_II_). Figure S1A shows that, for citrate buffer, the H_II_ is the only phase occurring at pH 5.5 at the three ionic strengths. In the case of acetate (Fig. S1B) Fd3m and H_II_ phases coexist at all ionic strengths, but an increase in ionic strength results in an increase in the signal of the inverse hexagonal structure compared to the Fd3m phase. The observed results in Fig. S1 can be ascribed to electrostatic interactions that play a role in buffer specificity. An increase of ionic strength has been found to result in a shift in the equilibrium of -NH^+^ ⇔ -N + H^+^ for triethanolamine buffer towards the left, corresponding to a shift to higher effective pH (that is lower H^+^ activity).^41^ Similarly, in our system at a fixed pH 5.5, an increase of ionic strength would result in a higher [-NH^+^]/[-N] ratio for the MC3 headgroup which would stabilize H_II_ with respect to the Fd3m phase. A possible mechanism for citrate is that the divalent buffer ions interact with MC3 positively charged MC3 headgroups, stabilizing the formation of the inverse hexagonal H_II_ phase. Nevertheless, the coexistence of the Fd3m and H_II_ phases, observed for acetate at the three ionic strengths, suggests that buffer specificity cannot be ascribed to a pure ionic strength (electrostatic) effect. Other mechanisms must also be at work.

We also investigated the effect of temperature, at 22°C and 37°C and pH 5.5. This is important to see if the transitions of the inner bulk phase of LNPs, which we are considering as the driving mechanism of mRNA transfection, occur in the same pH range observed above at body temperature. In Fig.S1B, we find the temperature increase favors the phase transition from inverse hexagonal H_II_ to Fd3m + H_II_ in the case of citrate and Fd3m + H_II_ to Fd3m for acetate. That is, the temperature increase has the opposite effect to an increase in ionic strength (Fig.S1A), tending to stabilize F3dm, that is, shifting the Fd3m - H_II_ transition to lower pH values. The temperature dependence is consistent with the trend we would expect from the van’t Hoff equation for the temperature dependence of the acid equilibrium p*K_a_*. From studies of the protonation enthalpy of ionizable alkylammonium ions^42^ we would expect the dissociation enthalpy for MC3 to have a value around ΔH ≈ +50 kJ/mol. The positive enthalpy means pK*_a_* should decrease as temperature increases, and this trend was observed experimentally in simpler ionizable alkylammonium ions. A lower p*K_a_* value means the -NH^+^⇔ -N + H^+^ equilibrium is pushed towards dissociation at higher temperatures, leaving the MC3 molecule uncharged. This can be interpreted as equivalent to pushing the system towards behavior that would be expected at higher pH (at lower temperatures). Hence an increase in temperature has the effect of pushing the phase transition to lower pH. In summary, the increase of both ionic strength and temperature result in opposite behaviors (Fig.S1). The effect of increasing ionic strength is to favor the inverse hexagonal phase, while that of increasing temperature is to favor the inverse micellar phases.

### Mechanism of pH transition and buffer specificity

In the following we focus first on the general pH-dependent L_II_-H_II_ transition in lipid mesophases. Transfection efficiency measurements (Fig. 2) showed a buffer specific effect that follows a (conventional) Hofmeister series, with citrate > phosphate > acetate. In SAXS measurements this ordering is also observed for the pH value of the bulk phase transition Fd3m-H_II_. The bulk phase transitions observed by changing pH and ionic strength involve the MC3 charge, but crucially also involve a change in the area per MC3 headgroup. The balance between these two properties determines the phase transition pH. Then, buffer specificity introduces shifts to that transition pH via specific adsorption of buffer ions at the water-lipid interface (Fig. 4A). Based on observations from molecular dynamics simulations as shown below, we find that the area of the MC3 headgroup plays a crucial role in determining the stable phase. MD simulations show (Fig. 4B) that the inverse micelle phase exhibits an area per headgroup of 0.53 nm^2^, while the inverse hexagonal phase is found with a smaller area per headgroup (0.45 nm^2^). The formation of the charge on MC3 at low pH stabilizes the H_II_ phase because of the higher charge per unit area that it obtains because of the lower area per headgroup. But at the same time, the area per headgroup affects the L_II_-H_II_ transition via the elastic bending energy of the lipid layer.^28^

**Figure 4.**
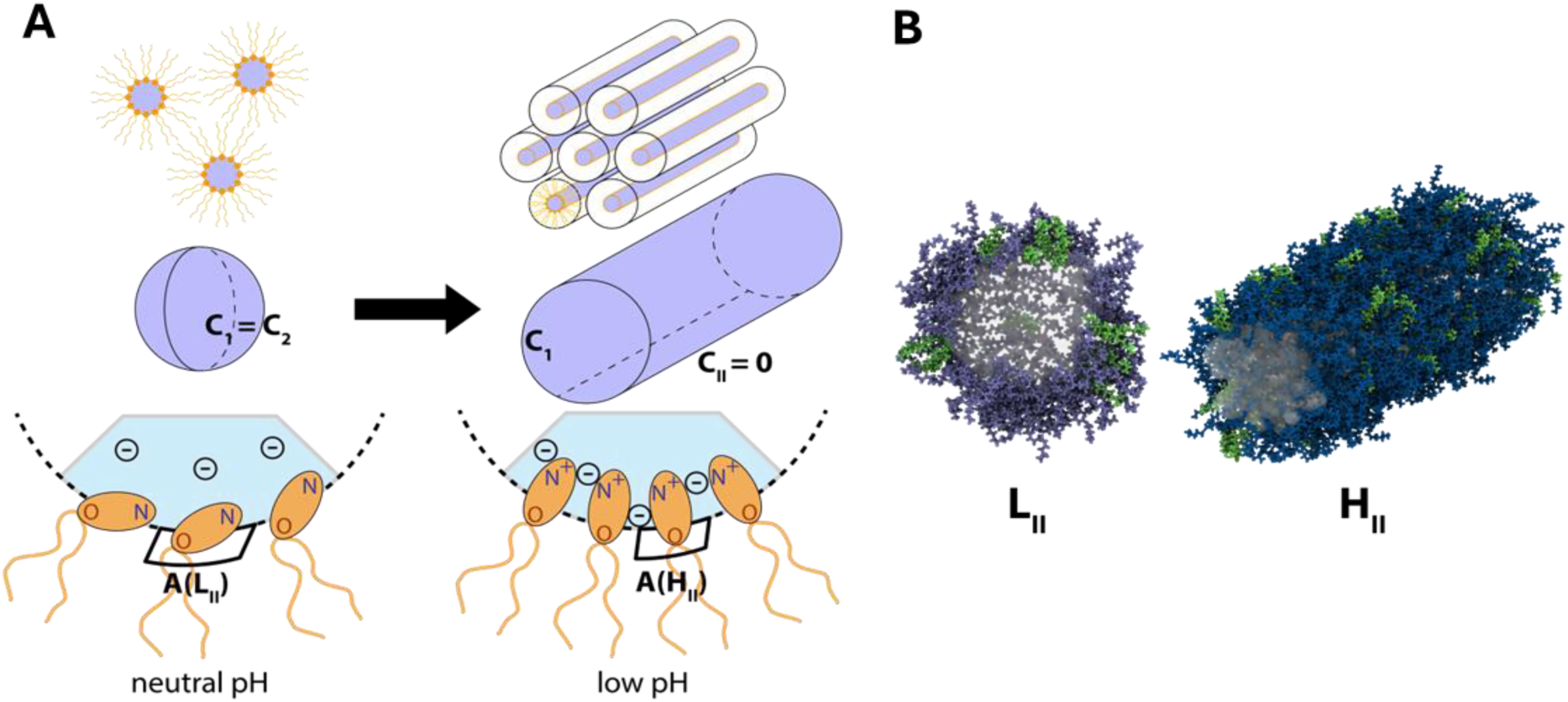
(A) Buffer effect on the curvature depending on pH. (B) MD simulations were performed for the inverse micellar (L_II_) phase with fully uncharged MC3 and the inverse hexagonal (H_II_) with fully charged MC3 lipids. The area per lipid for the L_II_ and H_II_ lipid systems obtained from MD simulations as A(L_II_)=0.53 nm^2^ and A(H_II_)=0.45 nm^2^ along with a snapshot of the corresponding molecular structure is provided. The protonated and neutral MC3 lipids are represented by dark and light blue colors respectively, cholesterol is represented by green and water by the white cloud.

The bending energy describes the elastic energy associated with the curvature of the lipid layer and driven by the freedom lipid molecules compress against each other in the layer. The strength of the bending energy is characterized by bending moduli κ and 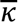, and may be quantified via a Helfrich bending energy *F_bend_ = 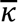(C_1_+C_2_)*^2^*/2+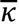C_1_C_2_. C_1_=1/R_1_* and *C*_2_=1/*R*_2_ are the curvatures associated with the radii *R_1_* and *R*_2_ in perpendicular directions along the surface of the lipid layer. Although spherical inverse micelles with *C*_1_=*C*_2_ have lower individual curvatures (larger radii, corresponding to larger d_NN_ values), the cylinders of the inverse hexagonal phase have a lower *total* curvature (*C*_1_+*C*_2_) because of the flat dimension along the axis of the cylinders with *C*_2_=0. In general, the two bending moduli κ, 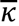 are distinct quantities. However, by applying Canham’s approximation,^43,44^ we may take the coarse estimate 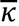 = −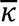. Under this coarse approximation the bending energies reduce to *F*_bend_[L_II_] ≈ 3κ*C*^2^ for generic inverse micelle phases, and *F* [H] ≈ κC_1_^2^/2 for the inverse hexagonal phase. In this case, for the same bending modulus the inverse hexagonal phase would be favored (lower energy), and the inverse micelle phases would not be observed. But at the same time, the bending modulus depends strongly on area per headgroup, κ∼*A*^-7.45^ The lower area per headgroup of the inverse hexagonal phase therefore means it has a far stronger bending modulus than the inverse micelle phases. Consequently, the bending energy favors the formation of inverse micelle phases with a larger area per headgroup. Thus, the higher surface charge density of the inverse hexagonal phase favors formation of H_II_ at low pH, while the higher area per headgroup (hence lower bending energy) of the inverse micelle phases favors them at high pH. The balance between the two explains the broad trend of the MC3 phase transitions, with the transition point lying at a pH close to the MC3 pK*_a_* (= 6.4). Here we have focused on the broad transition between inverse micelle phases collectively and the inverse hexagonal phase. It is likely that the distinct identity of 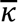, moving beyond Canham’s approximation, will be involved in transitions between specific inverse micelle phases such as P6_3_/mmc to F3dm or L_II_ to P6_3_/mmc.

To explain the buffer specific order observed for bulk phase transitions, we propose a mechanism of specific ion adsorption with a consequent buffer-specific change in the area per headgroup. As illustrated in Fig. 4A, we identify near (charged N group) and far (O ester group) moieties within the headgroup. The N-moiety remains in contact with the aqueous phase whereas the O-moiety may not be in contact, depending on the headgroup orientation (Fig. 4A). The strength of interactions of buffer ions with the lipid surface, including the N-moiety, follows the conventional series citrate > HPO_4_^2-^ > acetate, confirmed by our MD simulations (Fig. 5B). On the other hand, MD simulations (Fig. 5C), and evaluation of London dispersion coefficients of ions (Table S4) with the O-moiety, suggest that adsorption at the O-moiety would be stronger for acetate than other buffer ions. Adsorption of acetate at the O-moiety requires the ion to penetrate deeper into the lipid phase and therefore is expected to lead to an increase in the area per headgroup. In this way acetate stabilizes L_II_, consistent with SAXS data. At the same time, adsorption of negatively charged ions at the N-moiety is expected to reduce the repulsion between the positively charged MC3 headgroup and will therefore reduce the area per headgroup at low pH. Consequently, citrate anions with preferred adsorption at the (positively charged) N-moiety (Fig. 5B and Table 1) decrease the area per head group, increasing the bending modulus and shifting the Fd3m-H_II_ transition to a higher pH by stabilizing H_II_. Acetate ions with preferred adsorption at the O-moiety, likely involving London dispersion forces, increase the area per head group, decreasing the bending modulus and shifting the Fd3m-H_II_ transition to a lower pH by stabilizing Fd3m. This mechanism, involving two types of specific interactions between the buffer ions and the lipid headgroups, in particular with the positively charged N-moiety (via electrostatic interactions) and at the neutral O-moiety (via nonelectrostatic dispersion forces) is in agreement with the results obtained by investigating the effect of ionic strength (Fig. S1). In short, we can identify a direct buffer effect mediated via electrostatics and ion dispersion interactions, and an indirect effect via a consequent change in the area per head group.

**Figure 5.**
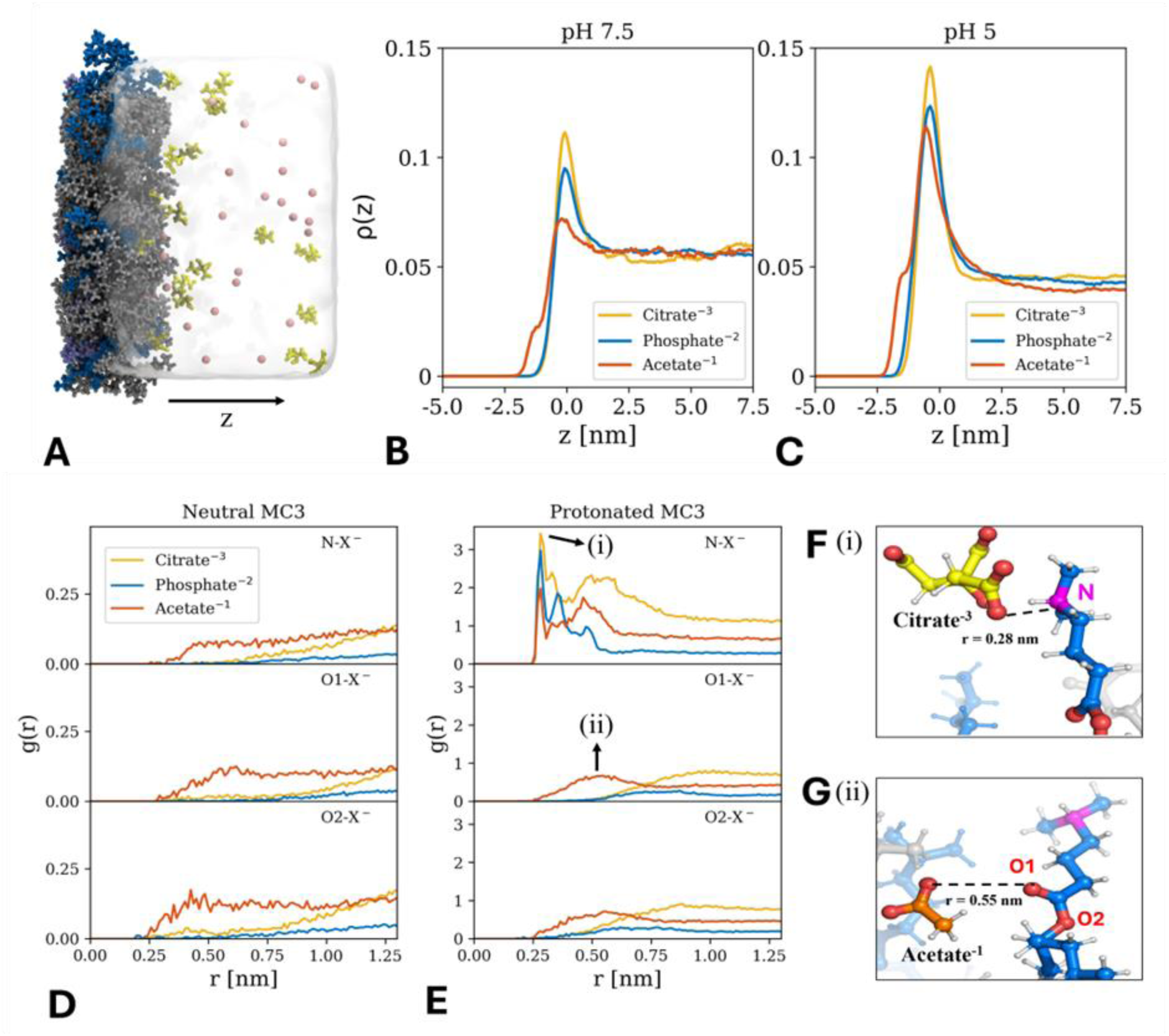
(A) Simulation snapshot of the lipid/water interface of the monolayer system at pH 5. Charged and uncharged MC3 lipids are shown in dark and light blue, respectively. POPC is shown in gray, citrate ions in yellow and sodium in pink. (B, C) Normalized probability distributions of the buffer ions for pH 7.5 and pH 5 along the z axis (perpendicular to the interface as indicated in A). (D, E) Radial distribution function g(r) of the different buffer ions around the nitrogen (N) and oxygen (O1, O2) atoms of an uncharged or charged MC3 molecule in the monolayer at pH 5. (F, G) Selected simulation snapshot of citrate and acetate at distances indicated by the arrows in panel E.

### All-atom MD simulations of interactions between buffer ions and MC3

To provide evidence of our hypothesis of specific ion adsorption at the lipid/water interface, we performed all-atom MD simulations in explicit water. To ensure the correct protonation degree of the ionizable MC3 lipid, we chose a MC3/POPC monolayer system (Fig. 5A) for which the area per lipid and protonation degree at pH 5 and 7.5 were determined consistently in previous work.^46^ In the simulations, citrate buffer was represented by the trivalent citrate ion, phosphate buffer by HPO_4_^2-^, and acetate buffer by the monovalent acetate ion. The affinity of the ions in terms of the probability distribution perpendicular to the lipid/water interface at pH 5 and 7.5 is shown in Fig. 5B, C. The main adsorption peak at z = 0 nm corresponds to the interactions of the ions with the nitrogen moiety and follows a conventional Hofmeister ordering: citrate > phosphate > acetate.

A second minor adsorption peak at z = −2 nm appears for acetate and corresponds to the interaction with the oxygen moiety. This peak is absent for the other ions and significantly smaller compared to the main adsorption peak. Further insights into the adsorption behavior at the interface can be gained from the local radial ion distributions *g*(*r*) of the around the charged and uncharged MC3 molecules. Fig. 5D, E shows *g*(*r*) for the buffer ions around the N and O-moieties (O1 representing the carbonyl O-moiety and O2 representing the ester O-moiety). For neutral MC3, the distributions at N- and O-moieties are similar (Fig. 5D). For charged MC3 (Fig. 5E), pronounced differences between the ions and the different binding sites can be observed. The results show that citrate ions have a higher affinity toward the N-atom in positively charged MC3 compared to the other buffer ions (as indicated by the highest peak in *g*(*r*) at a radial distance *r* = 0.28 nm in Fig. 5E, top). Phosphate ions have the second highest affinity for the N-moiety followed by acetate. In the case of O-moieties, only acetate ions adsorb while citrate and phosphate are depleted (Fig. 5E, bottom). The simulation snapshots reveal that citrate ions are located at the lipid/water interface while the acetate ions penetrate deeper into the lipid phase (Fig. 5 F,G). In summary, the affinity of buffer ions toward a lipid layer containing MC3 follows the Hofmeister series: citrate > phosphate > acetate. However, the local distribution of the ions is much more complex. The adsorption of acetate and depletion of phosphate and citrate at the oxygen are likely the cause of ion specific buffer effects.

## CONCLUSIONS

In this work mRNA-LNPs were prepared in three buffers, citrate, phosphate, and acetate. Transfection efficiency showed a dependence on the preparation buffer employed, in the order citrate > phosphate > acetate. To rationalize the buffer-specific efficiency we recall that the core phase of LNPs exhibits a dense ordered lipid phase, which is predominantly formed by cationic ionizable lipid MC3 together with cholesterol and hence is pH responsive. Note that DSPC and PEG-lipid are unlikely to partition into inverse phases. To explore the effect of buffer on pH response, we studied MC3-cholesterol bulk mesophases as core phase mimicking system using SAXS synchrotron measurements. We find that the critical phase transition from inverse cubic micellar to inverse hexagonal (Fd3m-H_II_) is dependent on the buffer in the order citrate > phosphate ∼ acetate and it is shifting by 1 pH unit in citrate (6.5-5.5) compared to acetate (5.5-4.5). In earlier work it was hypothesized that the Fd3m-H_II_ phase transition occurs in the excess lipid regions inside LNPs and plays a critical role in the pH-dependent endosomal fusion process.^8,14^ This view is consistent with our finding that the kinetics of gene delivery, specifically the onset of GFP transfection in single cell GFP expression time courses, is buffer specific and occurs in the same order as the critical pH values of the Fd3m-H_II_ transition. In the endosomes, which are slowly acidified, a larger critical pH value corresponds to an earlier transition.^47^ We assume that the structural transition of the LNP core induces endosomal fusion. This is likely as the inverse micellar-to inverse hexagonal transition causes defects that might propagate to the surface of LNPs. A second mechanism involved in the transition is the association of the protonated ionizable lipid MC3 with the surface layer of LNPs at low pH. A cationic surface charge, in turn, increases the likelihood of endosomal fusion. A central question in our study is whether we have a theoretical explanation for the shift in the pH value of the Fd3m-H_II_ transition as a function of the specific buffer used. We propose that competition of elastic bending energy and charging of the ionizable lipid head group leads to the predicted shift. A surprising insight was that the protonated MC3 lipid exhibits a smaller headgroup area due to a conformational change in the headgroup as seen by MD simulations. The dependence of the phase transition on the buffer is explained by specific ion adsorption at the nitrogen (N) and oxygen (O) moieties of the ionizable lipid, with a consequent change in the area per headgroup. This view is consistent with the finding that increasing the ionic strength stabilizes the H_II_ phase, likely favoring an even smaller headgroup area. By contrast, increasing the temperature to 37°C stabilizes the Fd3m phase due to a decrease in the pK*_a_* of MC3, which favors the neutral lipid species, pushing the phase transitions toward lower pH. There is scope for further work to confirm our interpretation that the area per headgroup varies with buffer ion adsorption. It seems likely that acetate adsorption at the O-moiety, with an associated increase in area per headgroup, is responsible for the swelling trend observed in the H_II_ phase with the nearest neighbor distance (*d_NN_*) increasing with pH, while decreasing in the case of citrate and phosphate buffers.

We have shown that the pH dependent structural transition in ionizable lipid/cholesterol phases is buffer-specific. The assumption that a similar structural transition occurs in the excess lipid regions of the LNP core phase would explain the buffer-specific timing of gene expression onset, if the transition of critical for endosomal fusion. For the argument to be conclusive we must assume that the buffer ions used in the preparation of LNPs remain inside the LNPs and are not washed out. Since diffusion of ions across lipid membranes is slow, there is good reason to believe that buffer species indeed remain trapped in LNPs. Note also that the degree of encapsulation and the size of LNPs is independent of buffer used in the formulation. To what extent the structure-activity relation for MC3 can be transferred to other ionizable lipids like KC2 or DLin remains to be seen. In recent work it has been shown that the two mainstream COVID-19 vaccine ionizable ALC-0315 and SM-102 lipids, show the same order of core phase changes as presented here.^8^ Our measurement of the buffer-specificity of the L_II_-H_II_ transition in ALC-0315/cholesterol and SM-102/cholesterol bulk phases is in agreement with an enhanced transfection efficiency in-vitro for LNPs prepared in citrate buffer. The proposed mechanism involves a direct buffer effect via specific adsorption of buffers, but also an indirect effect via changes in the area per lipid headgroup consequent to buffer adsorption. A crucial finding is the role of buffer (acetate) adsorption at the O-moiety of the headgroup, distinct from interactions with the N-moiety carrying the headgroup charge. This suggests an avenue for engineering LNP behavior, manipulating the transition pH by functionalization of the headgroup beyond simply its ionizable character. Consistent with the notion of the lipid packing parameter,^11,48^ controlling the area per headgroup, and thereby the strength of the lipid layer bending energy, is key to controlling the transition. To achieve rational design strategies to exploit the described buffer specific effects theoretical models are required that explain the specific ion adsorption mechanism using MD simulation as well as an elastic continuum description of the mesophases. The consistent buffer specific effects on LNP behavior demonstrate that understanding ionizable lipid mesophase transitions is crucial for analyzing mRNA transfection efficiency and further advancement of lipid formulations for gene therapy.

## METHODS

### Materials

DLin-MC3-DMA(O-(Z,Z,Z,Z-heptatriaconta-6,9,26,29-tetraem-19-yl)-4-(N,N-dimethylamino) butanoate; (MC3, 99 %), ALC-315 ([(4-hydroxybutyl)azanediyl]di(hexane-6,1-diyl) bis(2-hexyldecanoate)) and SM-102 (9-Heptadecanyl 8-{(2-hydroxyethyl)[6-oxo-6-(undecyloxy)hexyl]amino}octanoate) were purchased by MedChemExpress. 1,2-distearoyl-sn-glycero-3-phosphocholine, (DSPC, 99%), 1,2-distearoyl-sn-glycero-3-phosphoethanolamine-N-[amino(polyethylene glycol) (DSPE-PEG2000, 99%), cholesterol, sodium citrate dihydrate (99 %), citric acid (99 %), monobasic sodium phosphate (99 %), disodium hydrogen phosphate (99 %), hydrochloric acid (37 %), sodium hydroxide (97 %), sodium acetate (99 %), acetic acid (99.8 %), and citric acid (99.5 %) were purchased by Avanti Polar lipid Sigma Aldrich.

### Preparation of eGFP-mRNA LNPs

ARCA eGFP (Enhanced Green Fluorescent Protein) mRNA (APExBIO) was encapsulated in an LNP of four lipid components (DLin-MC3-DMA, DSPC, Cholesterol, DMPE-PEG2000) in a 50:10:38.5:1.5 molar ratio. mRNA and buffer (pH 3) were prepared in aqueous solution (0.075 mg/mL) while the lipid components (1.11 mg/mL MC3, 0.27 mg/mL DSPC, 0.52 mg/mL cholesterol, 0.15 mg/mL DSPE-PEG2k) were dissolved in ethanol. Concentrations were chosen to reach a final LNP concentration of 0.05 mg/mL mRNA concentration with an N/P ratio of 4 and a final buffer concentration of 50 mM. Microfluidic mixing was carried out with the NanoAssemblr^TM^ Spark^TM^ (Precision NanoSystems) at a volume ratio of aqueous:organic 2:1 ratio. Following mixing, LNPs were incubated 20 min at room temperature. To remove residual ethanol and buffers from the solution, LNPs were transferred to Slide-A-Lyzer™ MINI dialysis cups 3.5 kDa molecular weight cut-off (ThermoFisher Scientific) and dialysed into water for 18 h at room temperature. Size distribution was measured using a DynaPro® NanoStar™ (Wyatt) DLS device. Encapsulation efficiency was assessed using the Quant-it^TM^ RiboGreen RNA dye (Invitrogen).

### LNPs transfection efficiency

Cells were cultured in RPMI 1640 Medium (ThermoFisher Scientific) supplemented with 10% (v/v) FBS (Fetal bovine Serum, ThermoFisher Scientific, #10270106), 5 mM HEPES (GibcoTM, Thermofisher Scientific, #15630080) and 1mM Na-Pyruvate (GibcoTM, Thermofisher Scientific, #11360070) at 37°C, 5% CO_2_. For live-cell imaging, microstructures were prepared to allow single-cell culture. Therefore, 6-channel µ-slides (ibidi) coated with cell-repellent PVA were treated with PLPP N-(4-[benzoyl]benzyl)-N,N,N-triethylammonium bromide) (enamine) in an agarose and calcium peroxide solution and selectively illuminated with UV-light (365 nm). Selective illumination was facilitated with a photomask patterned with 20 × 20µm squares and a 80µm spacer. After illumination, the channels were rinsed with water and 0.5M HCl. The resulting squares were coated with laminin by incubation in a 20µg/mL laminin (BioLamina) working solution in PBS for 1h at 37°C.

Application of cells in medium for 1h led to self-assembly in a single cell pattern as depicted in Fig. 2A. eGFP-mRNA-LNPs were diluted in RPMI medium supplemented with FCS to a final concentration of 1ng/µL and incubated for 1h at room temperature to allow formation of a protein corona. Subsequently, LNP solution was applied for 1h and washed afterwards with L15-medium without phenol red (ThermoFisher Scientific). Cells were transferred to the microscope (Nikon TI Eclipse) and imaged over 30h with image acquisition every 10 min. Image analysis was then performed using our in-house python-based software including segmentation and background correction based on Schwarzfischer et al.^49^ to generate fluorescence trajectories.

### Bulk phase preparation: dialysis steps

MC3/cholesterol bulk phases were prepared at a range of pH conditions via three dialysis steps. First, the MC3 and cholesterol were dissolved in ethanol and mixed in a molar ratio of 3:1 (MC3: cholesterol) to a total lipid concentration of 56.1 mg/mL (MC3 46.7 mg/mL and cholesterol 9.4 mg/mL). The mixture was put into a dialysis cup with a molecular weight cut off of 3.5 kDa. Samples were first dialyzed against a 50 mM citrate buffer (pH 3) containing ethanol (at a volume ratio of 3:1, buffer: ethanol) for 48 hours. In the second dialysis step, the sample was dialyzed against PBS (1 mM KH_2_PO_4_, 155 mM NaCl, 3 mM Na_2_HPO_4_ 0.7 H_2_O, pH 7.4) for 48 hours. In the third step, the sample was dialyzed against NaCl 150 mM and the buffer of choice (citrate, acetate or phosphate 50 mM at pH 3.5, 4.0, 4.5, 5.0, 5.5, 6.0, 6.5, 7.0) with the required final pH for 48 hours. The samples at different ionic strengths (*I*) were done at different acetate, phosphate and citrate concentrations to reach *I*= 160 mM, *I*=200 mM, *I*=300 mM including the background salt NaCl 150 mM. The supernatant was removed from the cup and the solid precipitates were extracted for characterization by SAXS.

### SAXS measurements

Synchrotron small-angle X-ray scattering (SAXS) was carried out at the P12 BioSAXS EMBL beamline, PETRA III, DESY (Hamburg, Germany). The beamline instrumentation has been described previously.^14^ Further SAXS experiments were performed using an internal instrument at LMU. The crystallographic space groups of the liquid crystalline phases were determined from relative peak positions. All the measurements at P12 were performed in quartz capillaries. The scattering data background was subtracted by measuring empty capillaries.

### Calculation of nearest neighbor distance *d_NN_*

The nearest neighbor distance was calculated for each mesophase using the peak positions with as their respective miller indices following the formula shown in Table S2 and S3 (see Supporting information).^14^

### All-atom molecular dynamics simulations

Ion specific adsorption simulation setup: To investigate the adsorption of acetate, phosphate and citrate at the lipid/water interface, we used a monolayer setup. The lipid monolayer contained MC3 and POPC lipids in a 1:4 molar ratio and was constructed using the Men-Gen web server^50^ following the published procedure of our previous work.^46^ Each leaflet of the monolayer contained 200 lipids, separated by a water column of approximately 15 nm. The reason for choosing this setup was that the protonation degree at two pH values was determined consistently from a combination of simulations and scattering experiments.^46^ This was crucial since the protonation degree is inherently difficult to predict since it depends on the local lipid environment and is not directly accessible from experiments. Specifically, the lipid monolayer at pH 5 has a protonation degree of 67.5% ionizable MC3. At pH 7.5 the protonation degree is 14.5%. Both values significantly deviate from simple theoretical predictions.

For each pH value, the monolayers were simulated at a buffer concentration of 50 mM. Sodium acetate (CH_3_COONa), sodium phosphate (Na_2_HPO_4_), and sodium citrate (Na_3_C_6_H_5_O_7_) were used as in the experiments. Thus, a total of six simulations of the lipid monolayer were performed, covering two pH levels and three different buffers. Each simulation was repeated three times to improve the sampling statics.

The AMBER Lipid 17 force field was used to describe the POPC lipids. For the cationic and neutral MC3 molecules, we used recently developed force fields, which closely reproduce the experimental structure of lipid layers and are compatible with the AMBER force field family.^51^ The TIP3P water model^52^ along with Mamatkulov–Schwierz^53^ force field parameters for Na^+^ ions were also used. The citrate ion was described using the force field parameters obtained from Wright^54^ while the acetate and phosphate ion force field parameters were obtained from Kashefolgheta.^55^ The combination rule for the anion-cation interactions were modified to avoid crystallization artifacts. The resulting force field parameters are available at “(https://git.rz.uni-augsburg.de/cbio-gitpub/force-fields-buffer-ions).”

All-atom molecular dynamic simulations were performed using the GROMACS^56^ package (v-2024). A gradient descent algorithm was used to minimize the energy of the system. The simulations were performed in the NVT ensemble to ensure the correct area per lipid.^51^ The temperature was maintained at 293.15 K using the velocity rescale thermostat with a time constant of 1.0 ps. The Lennard-Jones potential was cut off and shifted to zero at 1.2 nm. Short-ranged electrostatic interactions were cut-off at 1.2 nm and the Particle Mesh Ewald (PME) method was used to evaluate long-range electrostatics. Hydrogen bonds were constrained using the LINCS algorithm and a time step of 2 fs was used. Each production run was performed for 70 ns. The first 20 ns of the simulation were discarded to account for equilibration and the rest of the trajectory was analyzed using GROMACS inbuilt modules and MDAnalysis.^57^ The trajectories were visualized, and snapshots were generated using visual molecular dynamics (VMD).^58^

Area per headgroup of H_II_ and L_II_ phases: to obtain the area per headgroup, we used data from our previous work,^14^ where we simulated the inverse hexagonal (H_II_) and inverse micellar (L_II_) phases. The H_II_ phase consisted of fully protonated Dlin-MC3-DMA (MC3) lipids combined with cholesterol in a 3:1 molar ratio, maintaining a water-to-lipid ratio (*n_w_*) of 12 to match the experimental lattice spacing of 60 Å. The L_II_ phase consisted of fully uncharged MC3 lipids using also *n_w_* = 12. Details on the calculation of the area per headgroup are provided in the supporting information.

## Author Contributions

**C.C.**: Conceptualization; Formal analysis; Data curation; Investigation; Methodology; SAXS measurements; SAXS data acquisition; Validation; Visualization; Writing - original draft; Writing - review & editing. **J.P.**: Formal analysis; Data curation; Investigation; Methodology; SAXS measurements; SAXS data acquisition; Validation; Visualization; Writing - original draft. **J.M.**: Formal analysis; Data curation; Methodology; Investigation; Writing - review & editing. **Ak.Su**.: Validation; Formal analysis; Software; Methodology **C.E.B.** SAXS data acquisition **N.S.**: Validation; Formal analysis; Software; Methodology **D.F.P.**: Conceptualization; Formal analysis; Software; Investigation; Validation; Visualization; Writing - original draft, Writing - review & editing; Supervision. **A.S.**: Conceptualization; Formal analysis; Investigation; Methodology; Validation; Visualization; Funding acquisition; Writing - original draft, Writing - review & editing; Supervision. **J.R.**: Conceptualization; Formal analysis; Investigation; Methodology; Validation; Visualization; Resources; Funding acquisition; Writing - original draft, Writing - review & editing; Supervision. The manuscript was written through contributions of all authors. All authors have given approval to the final version of the manuscript.

## Notes

The authors declare no competing financial interest.

## ASSOCIATED CONTENT

### Supporting information

SAXS measurements on ionic strength and temperature effects in the presence of citrate and acetate, DLS and zeta potential data of LNPs, Miller indices of the measured lipid phases together with formula of the lattice constant, Molecular simulations methods including adjustment of combination rules for the force field parameters, force field parameters, and calculation of area per lipid. London dispersion coefficients of buffer ions in water and in non-polar media (.doc).

## ACKNOWLEDGMENT

C.C. thanks Mobilità Giovani Ricercatori (MGR) 2023 project from University of Cagliari (UniCA) and MIUR (PON-AIM Azione I.2–DD n. 407-27.02.2018, AIM1890410-2). AS thanks the Italian Center for Colloid and Surface Science (CSGI) and Fondazione di Sardegna - FdS 2022 (F73C23001600007) for financial support. NS thanks Ana Vila Verde for sharing the force field parameters, help on the partial charges and fruitful discussions. We thank Ekaterina Kostyurina for helping with sample preparation. Deutsches Elektronen-Synchrotron (DESY) P12 BioSAXS EMBL beamline PETRA III (Hamburg, Germany) is greatly acknowledged for synchrotron beamtime under the project’s proposal SAXS-1247 and SAXS-1261. JR and NS were supported by the German Federal Ministry of Education and Research through BMBF Project 05K18WMA and 05K18EZA within the framework of the Swedish–German research collaboration Röntgen-Ångström Cluster. DFP acknowledges financial support under the National Recovery and Resilience Plan (NRRP), Mission 4, Component 2, Investment 1.1, Call for tender No. 1409 published on 14.9.2022 by the Italian Ministry of University and Research (MUR), funded by the European Union – Next GenerationEU– Project Title Structure and flow dynamics of Concentrated AMphiphilic BIOmolecules (CAmBio): driving the change to eco-sustainable surfactant formulations – CUP F53D23008790001-Grant Assignment Decree No. n. 1386 adopted on 01/09/2023 by the Italian Ministry of Ministry of University and Research (MUR). We also acknowledge the CINECA award under the ISCRA initiative, for the availability of high-performance computing resources and support. The authors gratefully acknowledge the scientific support and HPC resources provided by the Erlangen National High Performance Computing Center (NHR@FAU) of the Friedrich-Alexander-Universität Erlangen-Nürnberg (FAU) under the NHR project b119ee.

## Supporting Information

**Figure S1.**
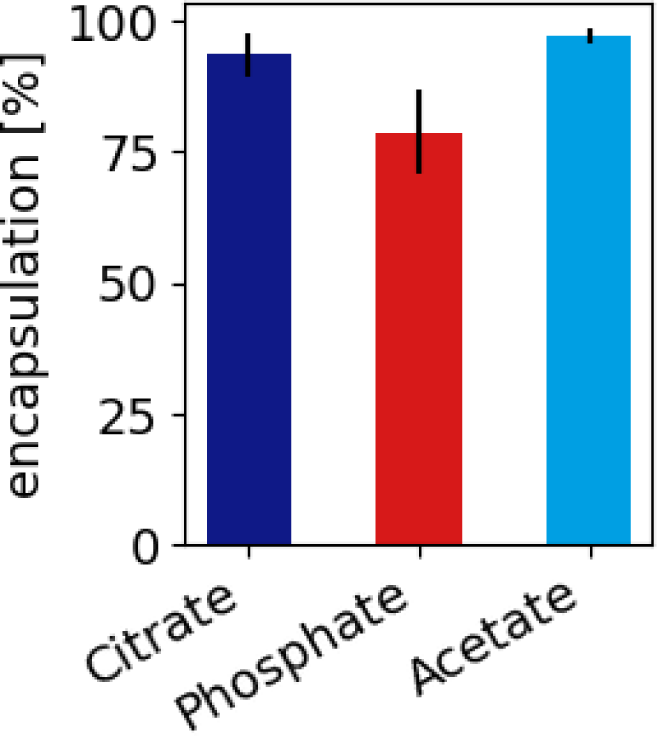
eGFP-mRNA LNPs encapsulation efficiency percentage in buffer citrate, phosphate and acetate 50 mM at pH 3. Encapsulation efficiencies are namely citrate 94 ± 4, phosphate 79 ± 8 and acetate 97 ± 2.

**Figure S2.**
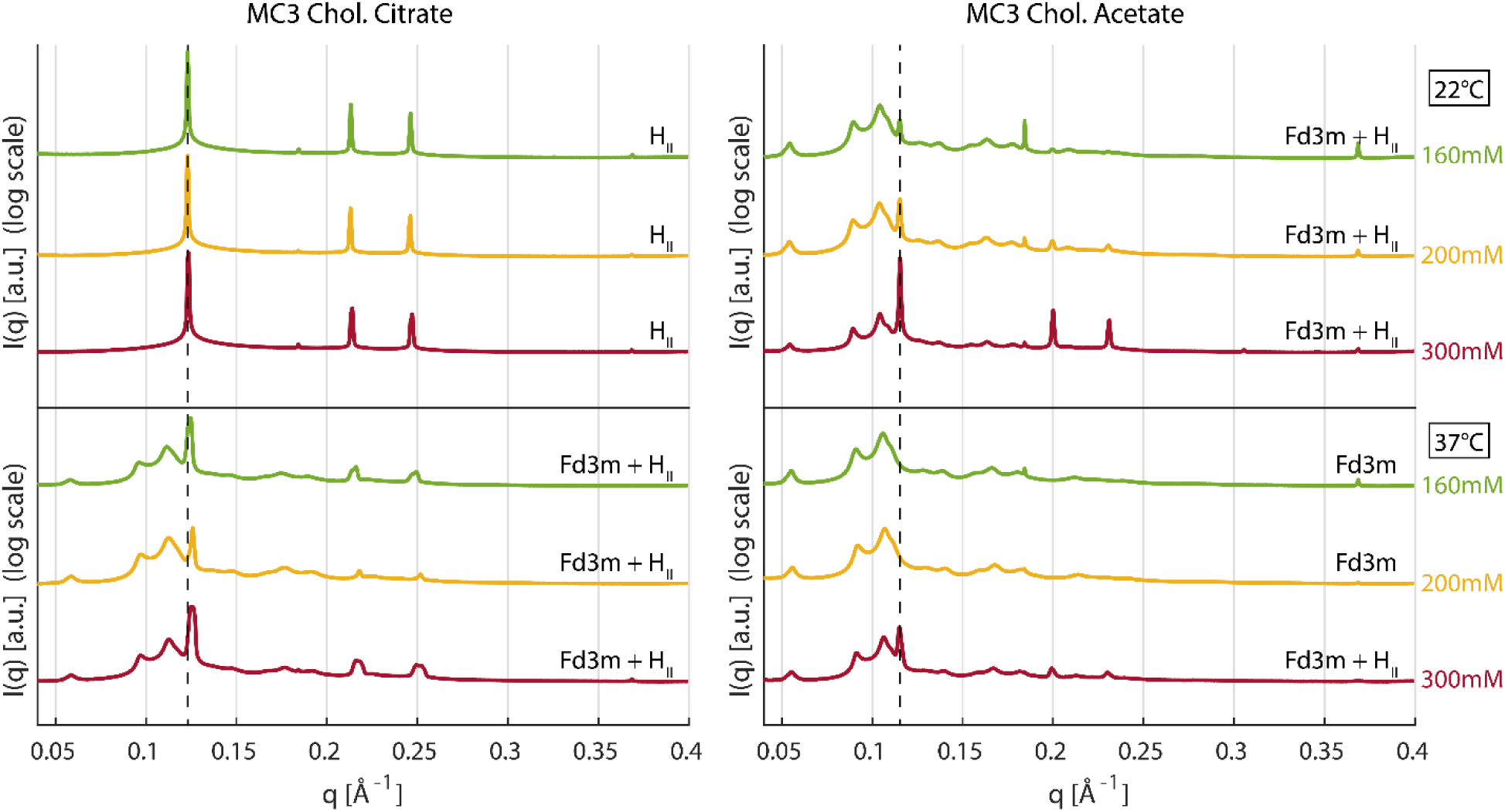
SAXS measurements of MC3-cholesterol dialyzed in the presence of 160-, 200-and 300-mM ionic strength citrate (A) and acetate (B) buffer at 22 °C and 37 °C at pH 5.5. Measurements of ionic strength (*I*) include the presence of NaCl 150 mM as background salt.

**Table S1.** Hydrodynamic diameter d_H_ a, Polydispersity index (PDI) as indication of the size measurements quality associated to the size distribution and zeta potential measurements of LNPs prepared in acetate 155 mM, phosphate 100 mM, and citrate 50 mM buffers at pH 4.5.

| Buffer | $d_H$ (nm) | PdI | pH* |
| --- | --- | --- | --- |
| Citrate | $76 \pm 7$ | 0.330 | $5.75 \pm 0.25$ |
| Phosphate | $70 \pm 1$ | 0.304 | $5.3 \pm 0.5$ |
| Acetate | $73 \pm 2$ | 0.324 | $5.25 \pm 0.25$ |

\* Fd3m- $H_{II}$ transition

**Table S2.**
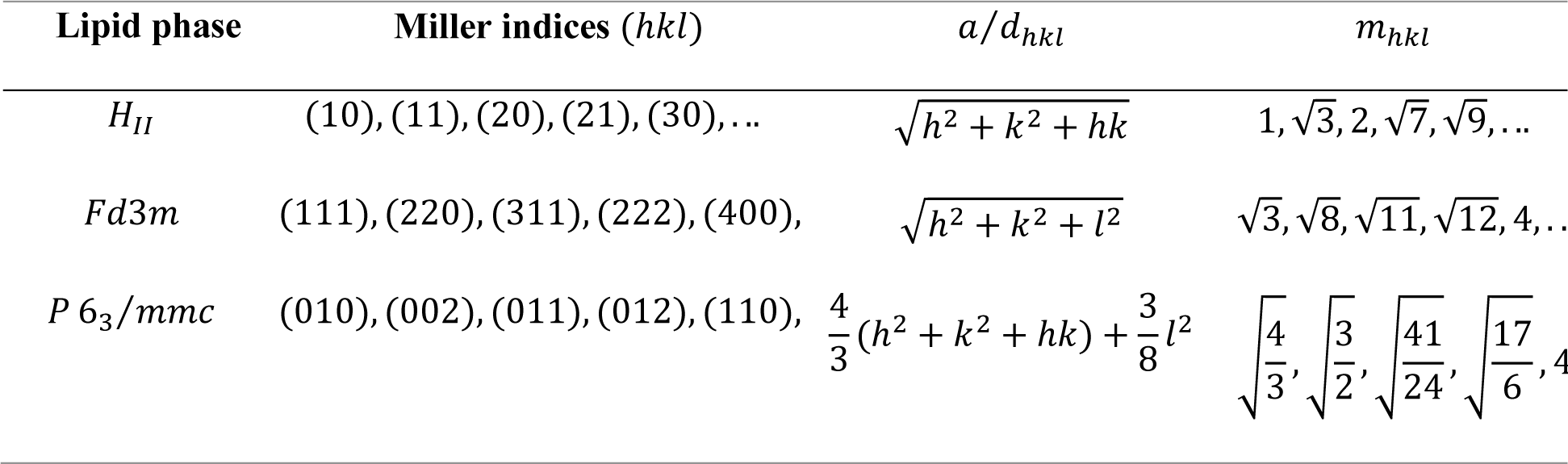
Miller indices of the measured lipid phases. *a* is the lattice parameter and *d*_*hkl*_ the corresponding real-world distance. This table is taken from reference ^1^.

| Lipid phase | Miller indices ($hkl$) | $a/d_{hkl}$ | $m_{hkl}$ |
| --- | --- | --- | --- |
| $H_{II}$ | (10), (11), (20), (21), (30), ... | $\sqrt{h^2 + k^2 + hk}$ | $1, \sqrt{3}, 2, \sqrt{7}, \sqrt{9}, \dots$ |
| $Fd3m$ | (111), (220), (311), (222), (400), | $\sqrt{h^2 + k^2 + l^2}$ | $\sqrt{3}, \sqrt{8}, \sqrt{11}, \sqrt{12}, 4, \dots$ |
| $P 6_3/mmc$ | (010), (002), (011), (012), (110), | $\frac{4}{3}(h^2 + k^2 + hk) + \frac{3}{8}l^2$ | $\sqrt{\frac{4}{3}}, \sqrt{\frac{3}{2}}, \sqrt{\frac{41}{24}}, \sqrt{\frac{17}{6}}, 4, \dots$ |

**Table S3.** Formula of the lattice constant *a* and the nearest neighbor distance *d*_*NN*_ for the measured lipid phases using the peak positions *q*_*hkl*_ as well as *m*_*hkl*_ from Table 1. This table is taken from reference 1.

| Lipid phase | $a$ | $d_{NN}$ |
| --- | --- | --- |
| $H_{II}$ | $\frac{4\pi m_{hk}}{\sqrt{3}q_{hk}}$ | $a$ |
| $Fd3m$ | $\frac{2\pi m_{hkl}}{q_{hkl}}$ | $\frac{a}{\sqrt{8}}$ |
| $P 6_3/mmc$ | $\frac{2\pi m_{hkl}}{q_{hkl}}$ | $a$ |

**Figure S3.**
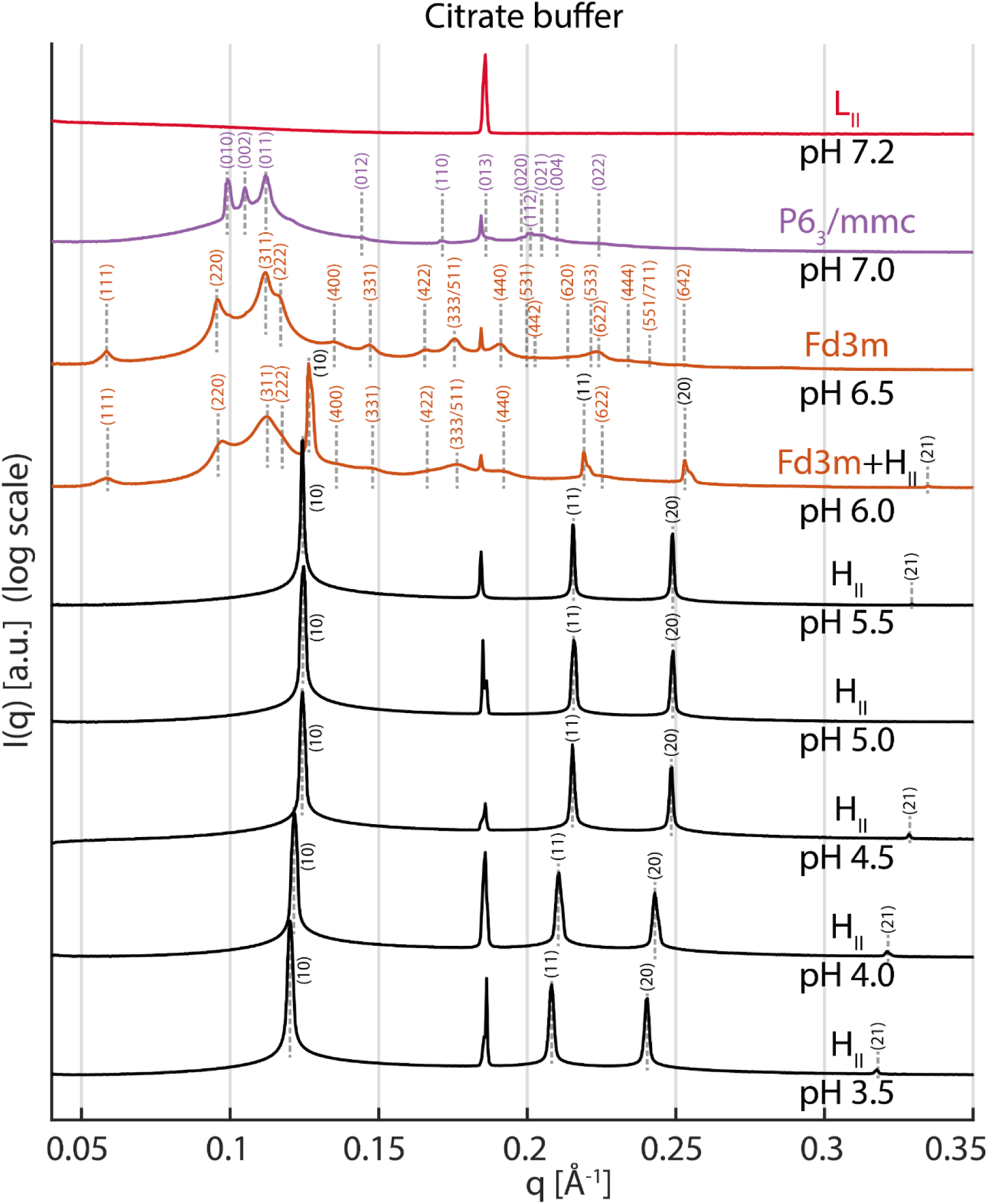
SAXS measurements and Miller indices of the mesophase of MC3-cholesterol samples dialyzed in the presence of 50 mM citrate buffer and NaCl 150 mM in a range of pH ≈ 3.5 - 7.5.

**Figure S4.**
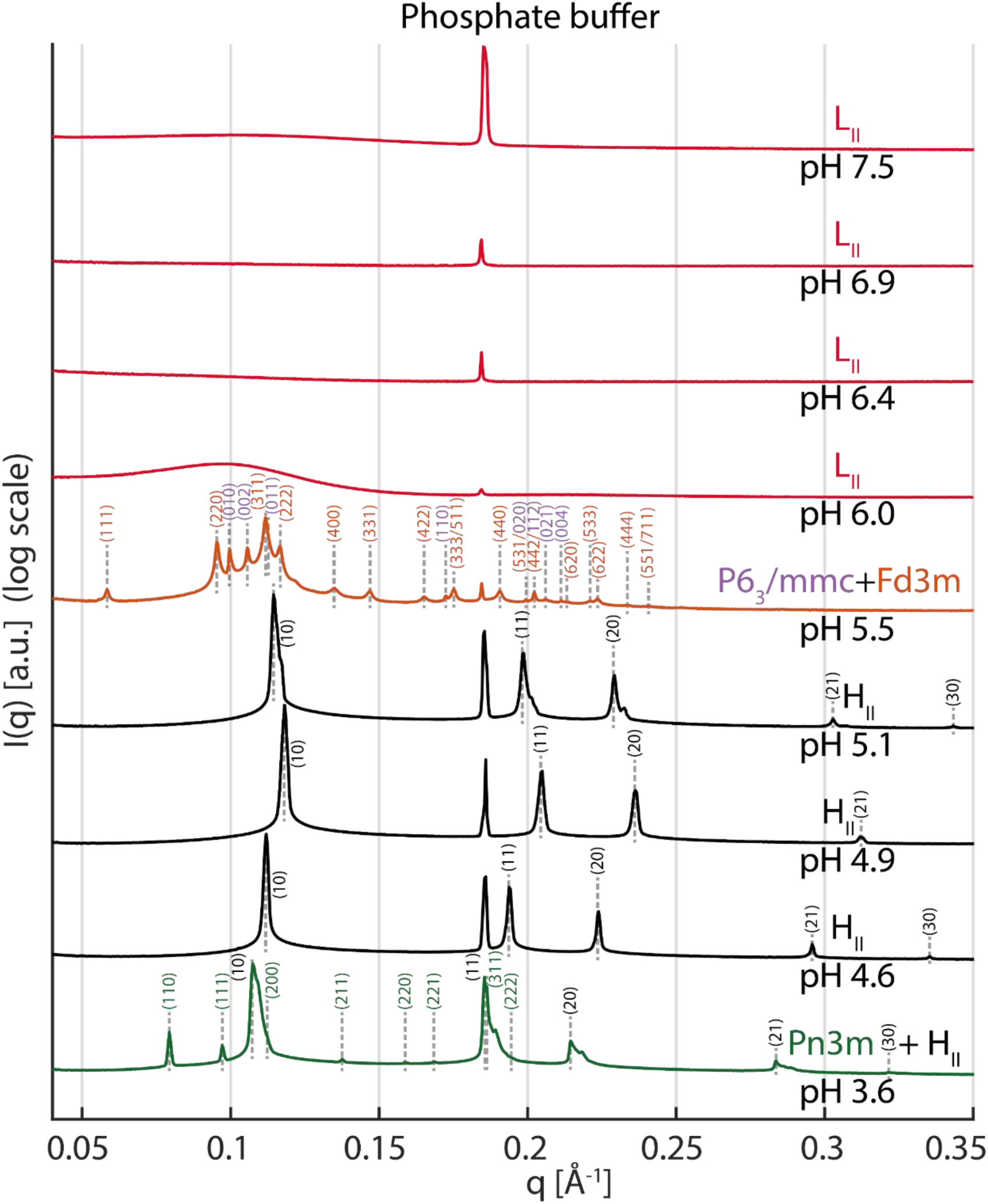
SAXS measurements and Miller indices of the mesophase of MC3-cholesterol samples dialyzed in the presence of 50 mM phosphate buffer and NaCl 150 mM in a range of pH ≈ 3.5 - 7.5.

**Figure S5.**
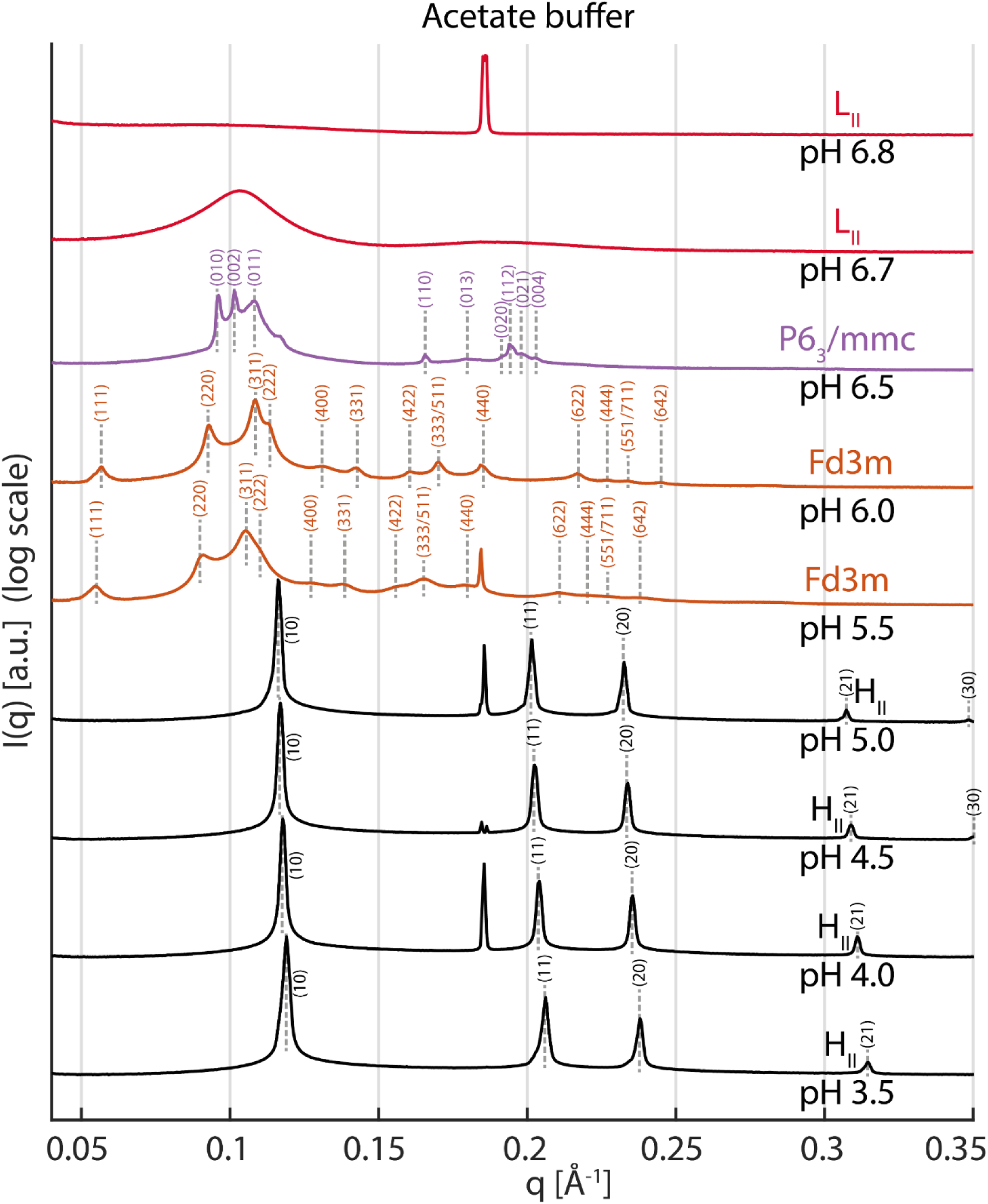
SAXS measurements and Miller indices of the mesophase of MC3-cholesterol samples dialyzed in the presence of 50 mM acetate buffer and NaCl 150 mM in a range of pH ≈ 3.5 - 7.5.

**Figure S6.**
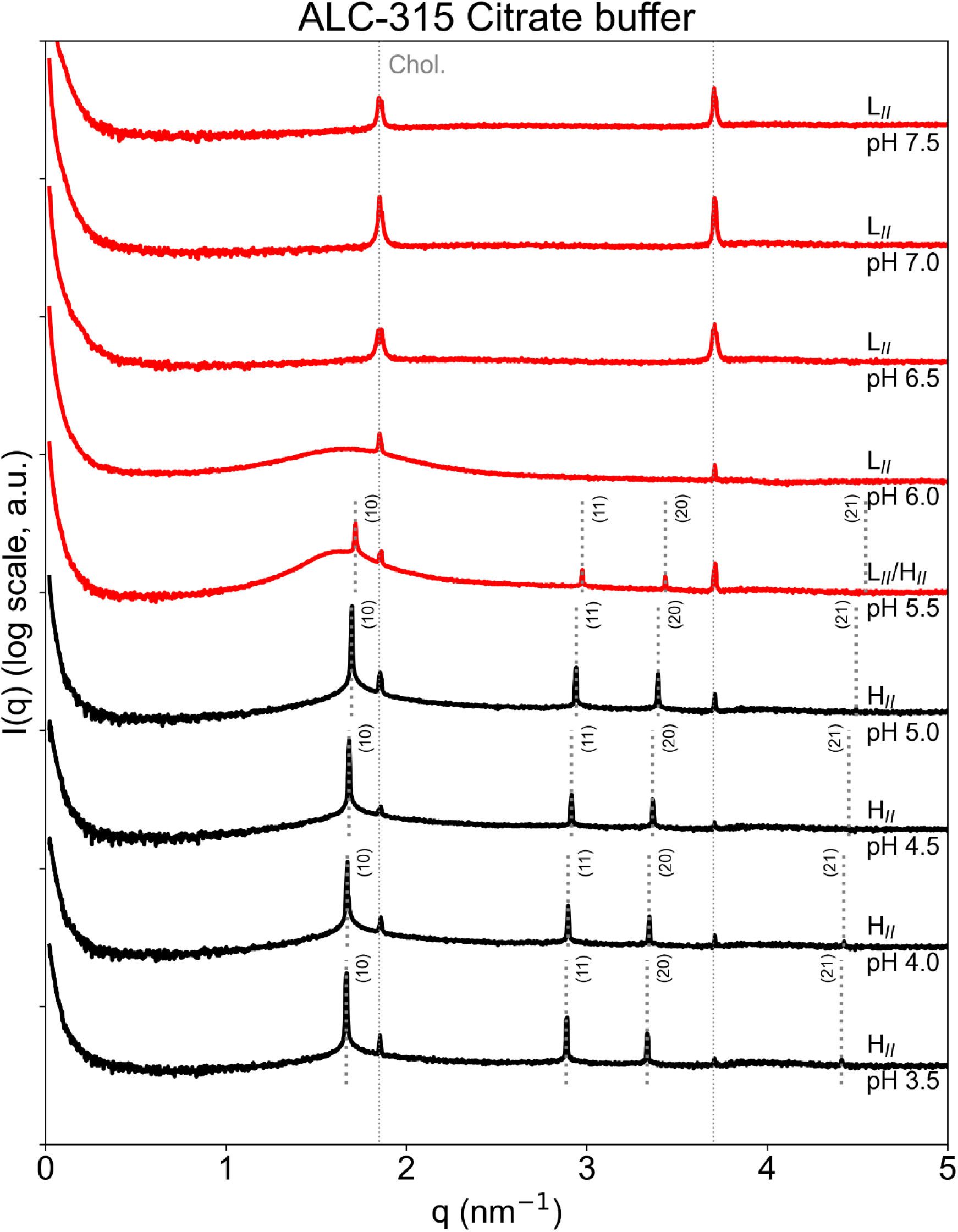
SAXS measurements of the mesophase of ALC315-cholesterol samples dialyzed in the presence of 50 mM citrate buffer and NaCl 150 mM in a range of pH ≈ 3.5-7.5.

**Figure S7.**
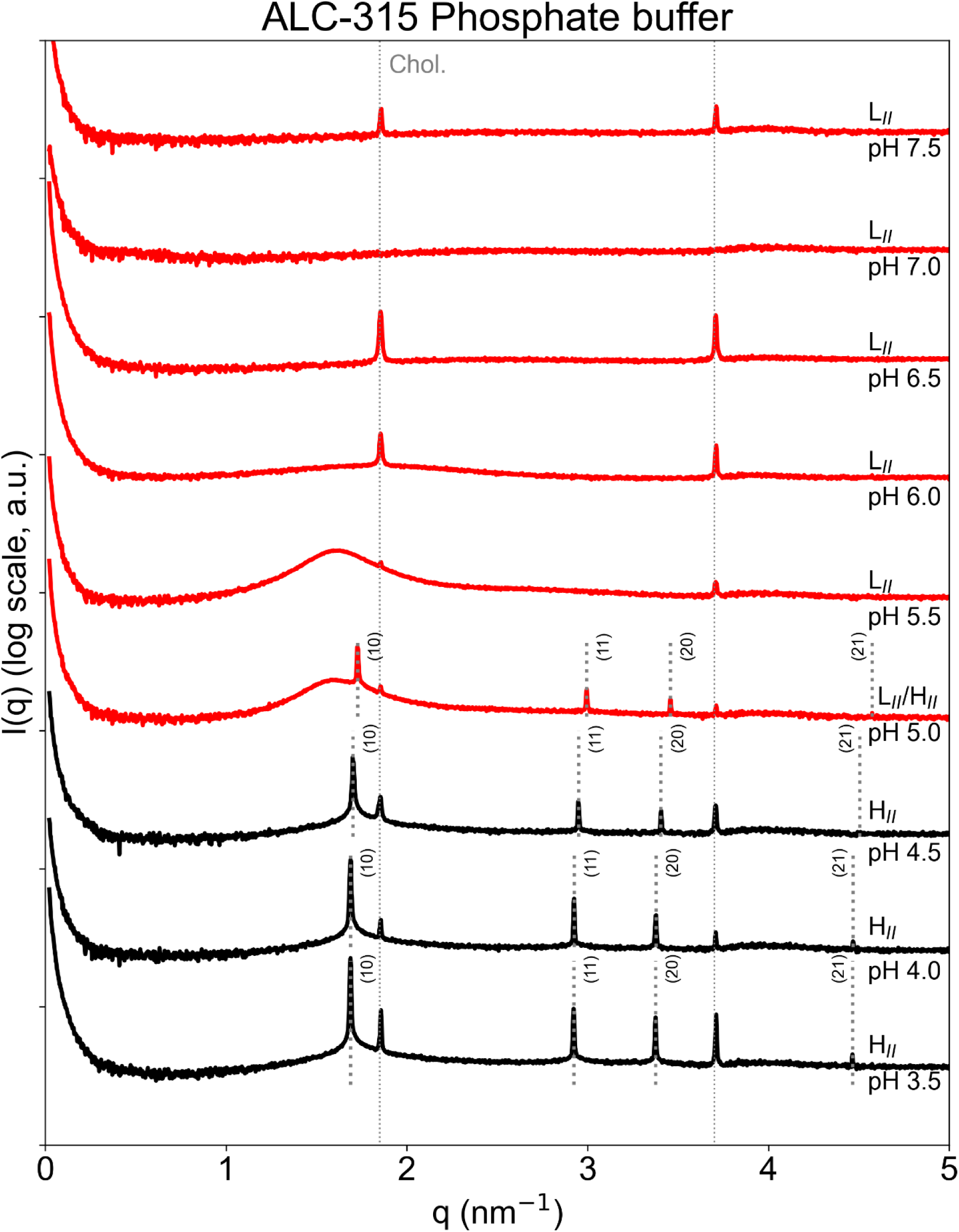
SAXS measurements of the mesophase of ALC315-cholesterol samples dialyzed in the presence of 50 mM phosphate buffer and NaCl 150 mM in a range of pH ≈ 3.5-7.5.

**Figure S8.**
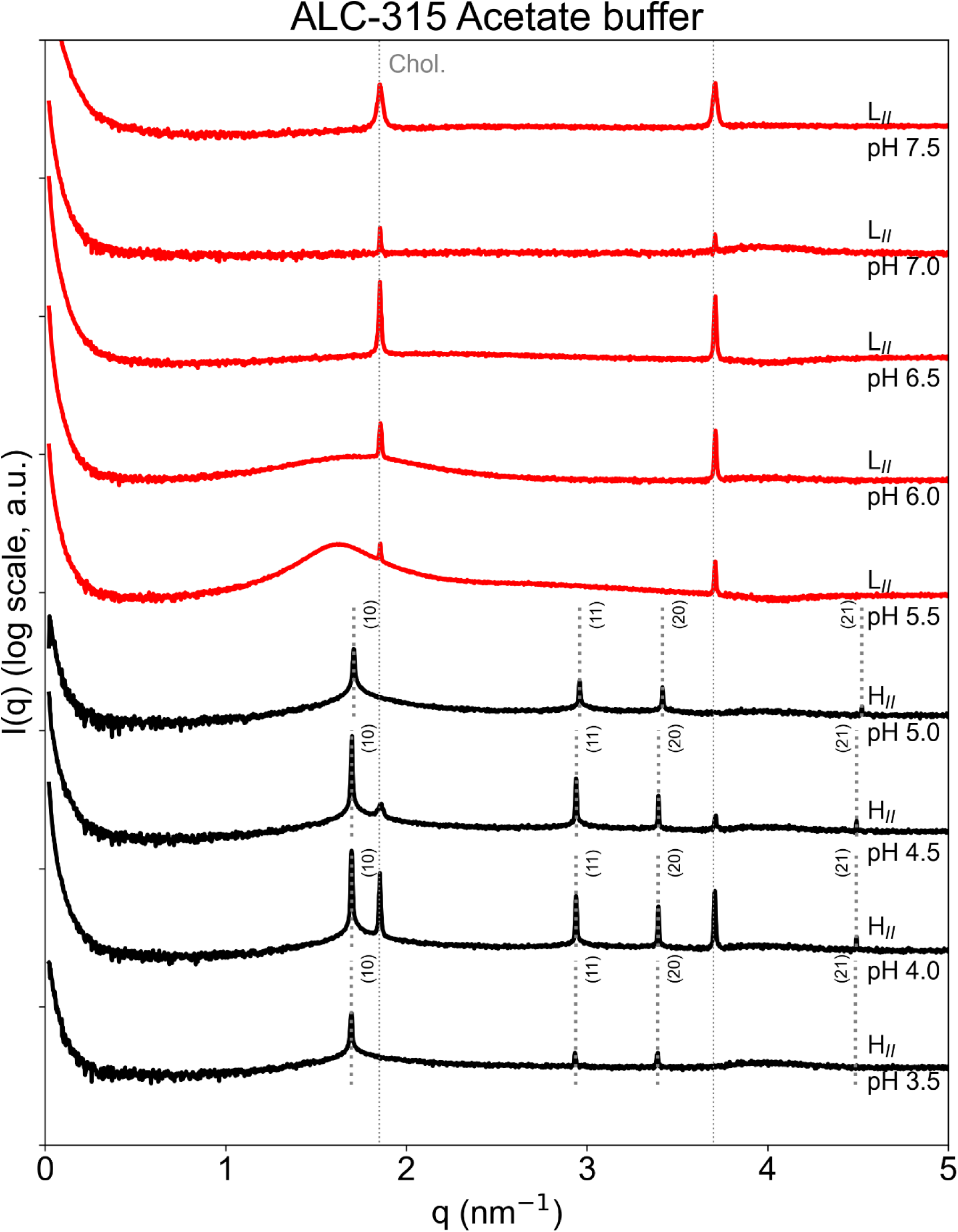
SAXS measurements of the mesophase of ALC315-cholesterol samples dialyzed in the presence of 50 mM acetate buffer and NaCl 150 mM in a range of pH ≈ 3.5-7.5.

**Figure S9.**
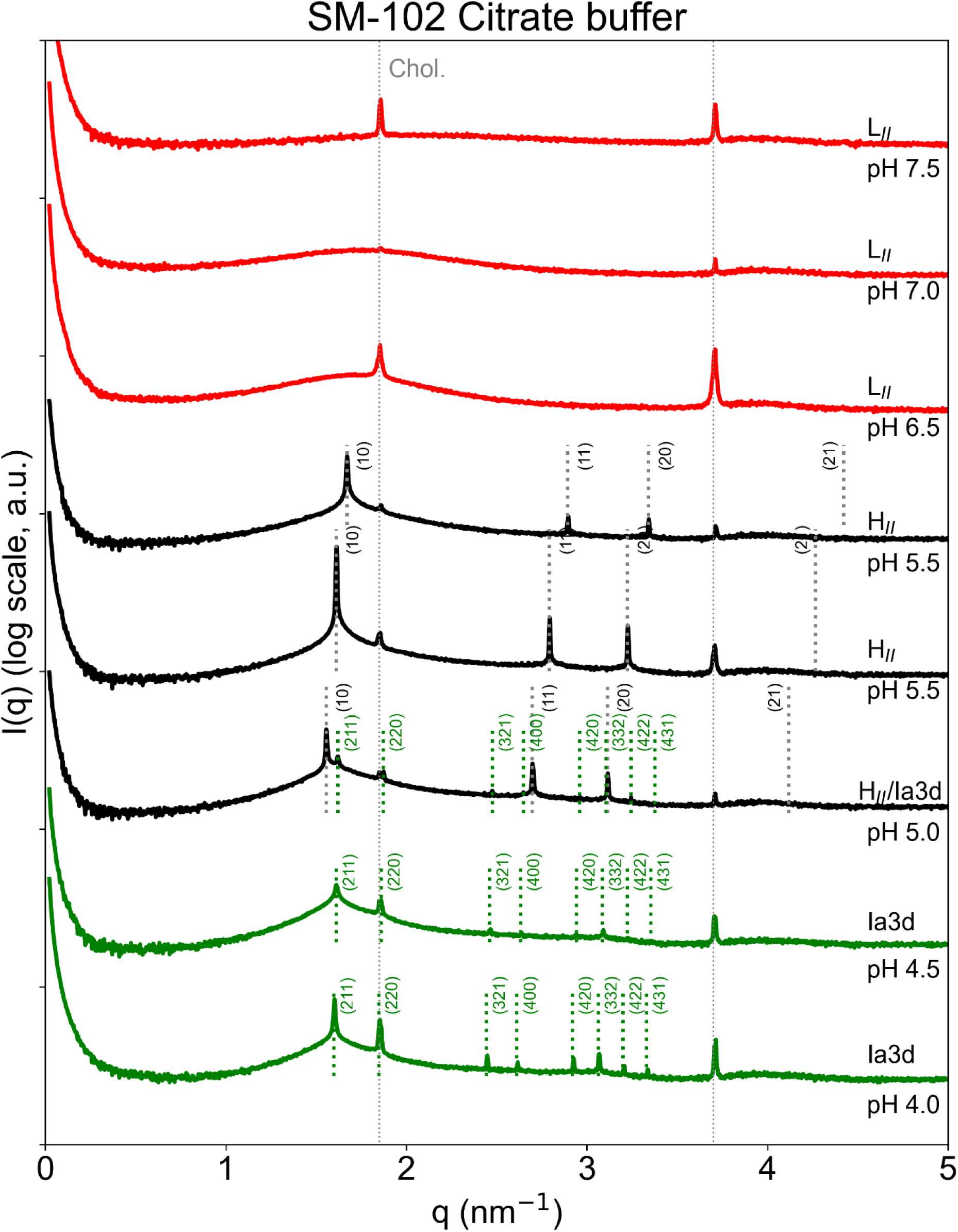
SAXS measurements of the mesophase of SM102-cholesterol samples dialyzed in the presence of 50 mM citrate buffer and NaCl 150 mM in a range of pH ≈ 4.0 - 7.5.

**Figure S10.**
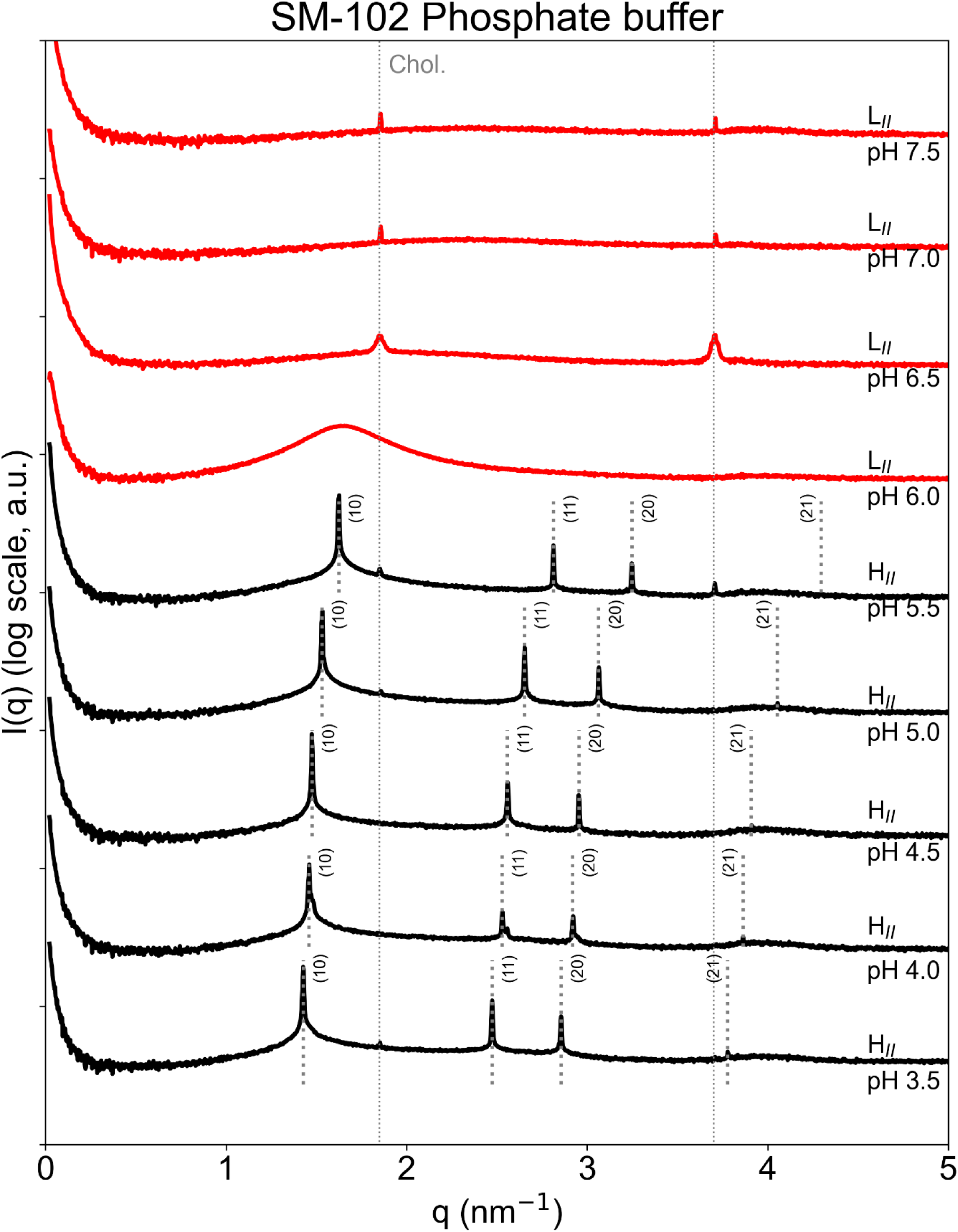
SAXS measurements of the mesophase of SM102-cholesterol samples dialyzed in the presence of 50 mM phosphate buffer and NaCl 150 mM in a range of pH ≈ 3.5 - 7.5.

**Figure S11.**
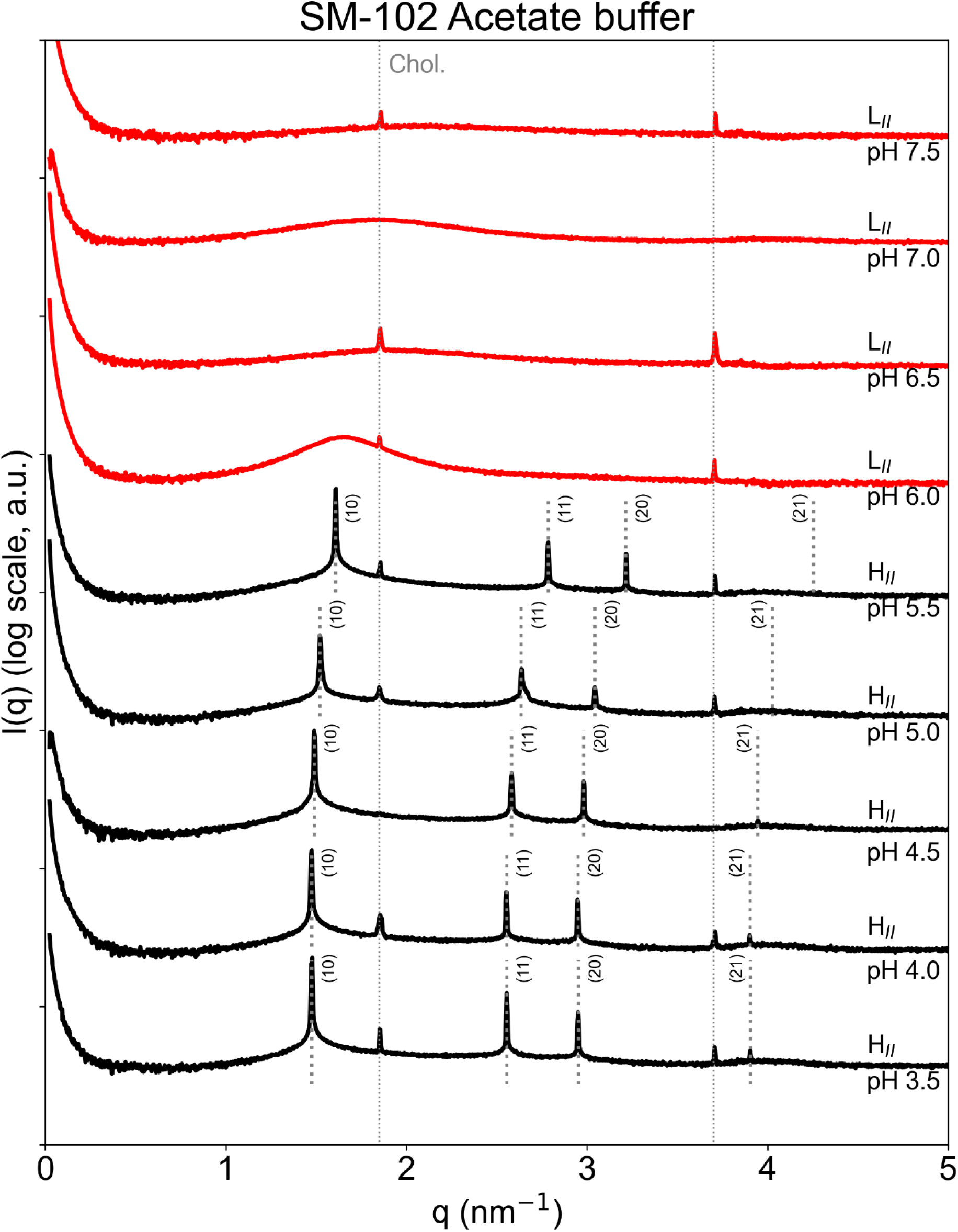
SAXS measurements of the mesophase of SM102-cholesterol samples dialyzed in the presence of 50 mM acetate buffer and NaCl 150 mM in a range of pH ≈ 3.5 - 7.5.

**Figure S12.**
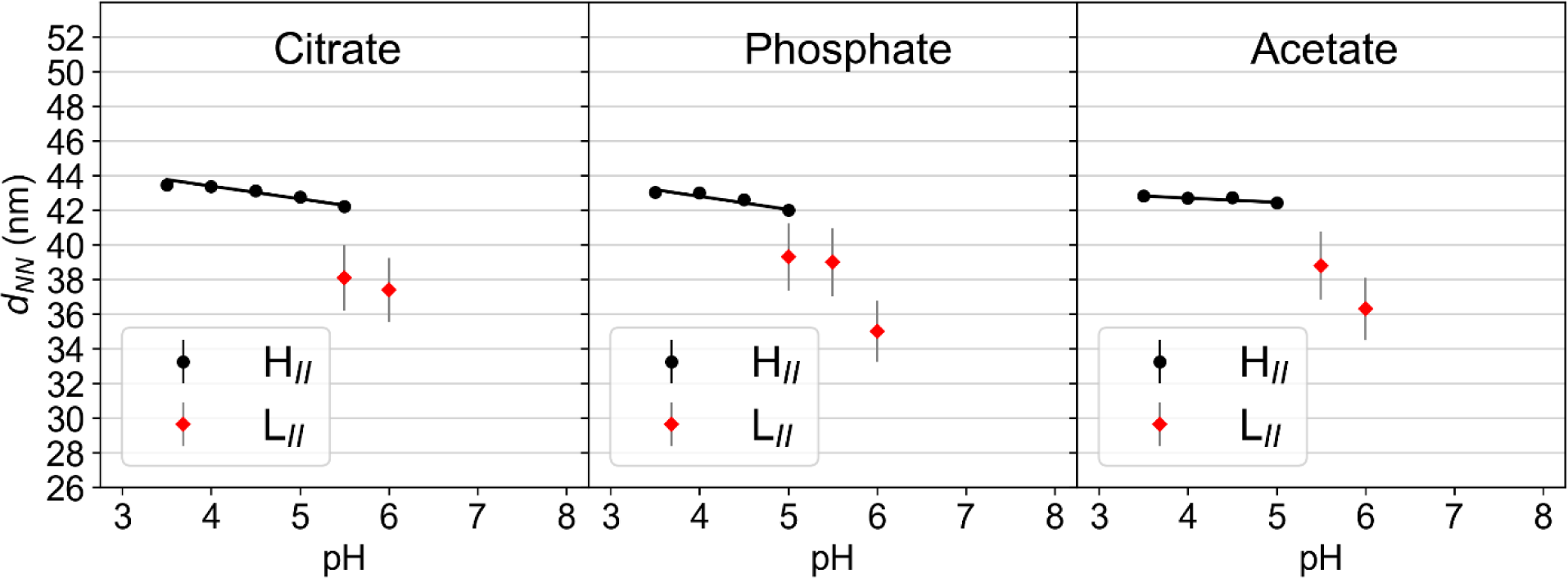
d_NN_ values for H_II_ and L_II_ phases for ALC315-cholesterol samples as a function of pH for buffer citrate, phosphate and acetate.

**Figure S13.**
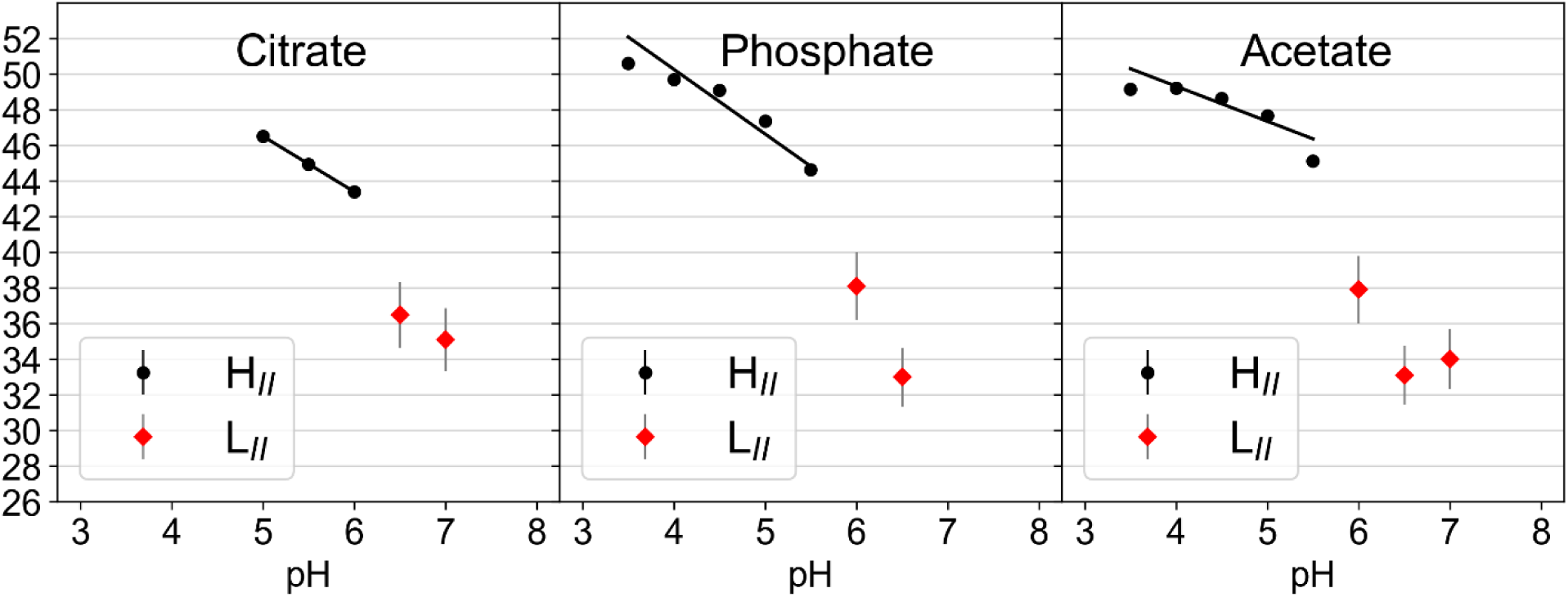
d_NN_ values for H_II_ and L_II_ phases for SM102-cholesterol samples as a function of pH for citrate, phosphate and acetate.

## Molecular simulation methods

### Adjustment of combination rules for the force field parameters

In MD simulations, the non-bonded interaction potential of the ions has the following form:

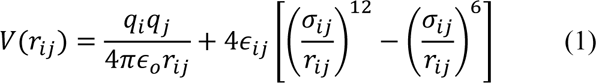

The first term describes the electrostatic interactions via a Coulomb potential. The last two terms describe the Van-der-Waal interactions via a 12-6 Lennard-Jones (LJ) interaction potential. *q_i_* and *q_j_* are the charges of atoms *i* and *j* respectively, *r_ij_* is the interatomic distance, *σ*_*ij*_ describes the LJ diameter at which the potential is zero and *ε*_*ij*_accounts for the interaction strength. MD simulations of citrate and phosphate ions at finite concentrations resulted in crystallization of the ions. This effect was observed for different Na^+^ force fields including the ones from ref. 35, 37 of the main manuscript. To avoid this artifact, we systematically increased the anion-cation LJ diameter by introducing a scaling factor in the corresponding Lorentz-Bertholot combination rule:

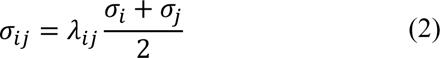

The value of *λ*_*ij*_ was increased until no more clustering was observed as judged by the anion-cation radial distribution functions. The final values are λ=2 for the citrate-sodium scaling and the phosphate-sodium interaction. Only the interaction of the sodium ions with the oxygen atoms of the phosphate and citrate ions were scaled. Note that in a more rigorous approach, the experimental activity derivative could be used to further optimize the anion-cation interactions as in previous work in reference 2.

### Force Field parameters for phosphate

The Lennard-Jones force field parameters for the phosphate ions (HPO_4_^2-^) were taken from H PO ^-^ in ref 3. The partial charges were obtained from RESP fitting.

### Calculation of area per lipid

The area per lipid of both the L_II_ and H_II_ phases were calculated for the system with n_w_ = 12. In the H_II_ phase, which is enclosed within a triclinic unit cell characterized by a lattice spacing *d_H_* and height *h*, we assume that the water column is an equally long cylinder with a radius *R_H_*. The water volume fraction (H) of the H_II_ phase is obtained by dividing the volume of the water column by the total volume of the system. The total volume of the system in the H_II_ phase is the volume of the triclinic unit cell which is given by *d_H_^2^h*sin(60°). The water volume fraction *ϕ*_*H*_ is given by:

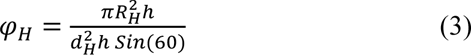

From this equation, the radius *R_H_* of the cylindrical water column can be derived as

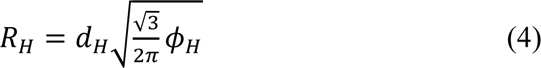

The value of H=0.24 and d_H_=60Å is taken from ref. 1. The area per lipid *(A)* of the f the H_II_ phase is obtained by dividing the surface area of the water column by the total number of lipids *N*.

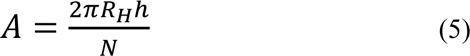

Similarly, for the L_II_ phase simulated in a rhombic dodecahedron box of lattice spacing *dL*, we assume a water sphere of radius *R_L_* surrounded by lipids. The water volume fraction (L) of the L_II_ phase is obtained by dividing the volume of the water sphere by the total volume of the system. For a rhombic dodecahedron box of length *dL*, the volume of the box is 0.707*d_L_*^3^ (i.e. *d*^3^/√2). The water volume fraction *ϕ*_*L*_ is given by

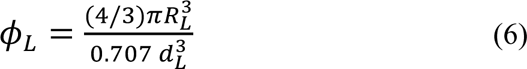

The radius of the water sphere *R_L_* can be obtained from the above equation by:

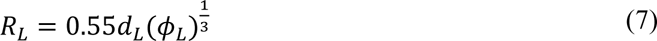

Corresponding area per lipid of the L_II_ phase,

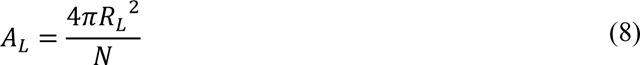

### Calculation of probability distributions

The probability distributions for the ions perpendicular to the interface were calculated as

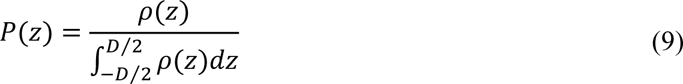

where *D* is the distance between the monolayers and ⍴(z) is the number density of the ions. The probability distributions are normalized such that 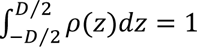.

### Calculation of radial distribution function

The radial distribution function *g*(*r*) provides insights into the local distribution of the ions around the protonated and neutral MC3 lipids. The radial distribution functions were obtained from the following equation:

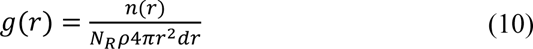

n(r) is the count of buffer ions around the reference atoms at distance *r*, *N_R_* is the number of reference atoms, and ⍴ = *N_I_*/*V_w_* is the bulk density of the ions. *N_I_* is the total number of anions in the system minus the anions used for neutralization and *V_w_* is the volume of water obtained by multiplying the theoretical molecular volume of a water molecule with the total number of water molecules in the system. In a bulk solution, *g*(*r*) gives the number of ions at distance *r* relative to the number of ions in an ideal solution. At an interface, such as the lipid water interface investigated here, equation (8) gives the number of buffer ions at distance r relative to a uniform ion distribution in the same slab geometry. The MD analysis python module (reference 4,5) was used to count the number of anions around the reference atoms (N, O1 and O2) of the charged and uncharged MC3 lipid head group using the simulation setup at pH 5.

### London dispersion interactions and the O-moiety

Acetate ion binding to the O-moiety of the MC3 headgroup can be understood in part from direct London dispersion interactions, *U*(*d*) = −*C*/*d*^6^, where *d* is the distance between the ion and the O-moiety. The London dispersion coefficient may be evaluated from quantum mechanical electronic polarizabilities of the species by applying quantum electrodynamics methods.6,7 Table S4 presents calculated London dispersion coefficients of buffer ions with a neutral acetic molecule representing the O-moiety (i.e. representing with O-moiety via a simple bound carbonyl group). London coefficients are evaluated in aqueous medium, and in nonpolar medium (tetradecane).

**Table S4.** London dispersion coefficients of buffer ions with acetic acid (representing the MC3 O-moiety) in aqueous media and in non-polar medium.

| Buffer ion | $C$, aqueous ($\text{\AA}^6 \text{ kJ/mol}$) | $C$, in tetradecane<br>($\text{\AA}^6 \text{ kJ/mol}$) |
| --- | --- | --- |
| acetate | 5414 | 3752 |
| $\text{H}_2\text{PO}_4^-$ | 5635 | 4473 |
| $\text{HPO}_4^{2-}$ | 4188 | 1722 |
| $\text{PO}_4^{3-}$ | -2069 | -8693 |
| $\text{H}_2\text{-citrate}^-$ | 2182 | 149 |
| $\text{H-citrate}^{2-}$ | 1456 | -1347 |
| $\text{citrate}^{3-}$ | -258 | -4635 |

The London coefficient is significantly more attractive for acetate ion than any of the citrate species. Curiously, trivalent citrate even has a negative coefficient, indicating a repulsive interaction pushing citrate away from the representative O-moiety. The aqueous London coefficient for phosphate is similar to acetate, indicating that this mechanism does not solely control the binding of acetate to the O-moiety observed in MD simulations. The O-moiety is located deeper inside the headgroup layer, such that the environment of the interaction is not purely aqueous. The reduced polar environment deep inside the head group is more unfavourable to phosphate than to acetate, which is surface active (partially oleophilic) due to its short hydrocarbon group. Indeed, the London coefficient of acetate in a nonpolar environment (tetradecane) is much stronger (more attractive) than all phosphate and citrate species apart from H_2_PO_4_^-^.

